# An ancient alkalinization factor informs *Arabidopsis* root development

**DOI:** 10.64898/2025.12.22.695669

**Authors:** Kaltra Xhelilaj, Michelle von Arx, David Biermann, Aleksander Parvanov, Natalie Faiss, Isabel Monte, Pascal Pireddu, Felix Klingelhuber, Cyril Zipfel, Marja Timmermans, Claudia Oecking, Julien Gronnier

## Abstract

The power of hydrogen (pH) regulates virtually all cellular activities. In both plants and animals, cell-to-cell variations in pH correlate with key developmental transitions^1–5^, yet the underlying regulators and associated functions remain elusive. Here, we report that members of the REMORIN (REM) protein family function as inhibitors of the H^+^-ATPases thereby promoting extracellular pH (pHe) alkalinization. This, in turn, regulates various cell surface processes, including steroid hormone signaling, and coordinates developmental transitions in the *Arabidopsis thaliana* root. Inhibition of H^+^-ATPases by REMs represents an evolutionary innovation that predates the origin of the root system itself. This study thus uncovers an ancient alkalinization mechanism co-opted by the root developmental program and infers that pHe patterning may have shaped morphogenesis evolution.

## Main

The concentration of protons (H^+^), also known as the power of hydrogen (pH), is of paramount importance for all forms of life. The pH modulates the physicochemical properties of biomolecules, thereby dictating a plethora of cellular activities^6,7^. Cells sustain and tightly regulate subcellular pH optimums that specify organelles function^8–10^. In addition, in multicellular organisms cell-to-cell variations in pH have been linked with key developmental transitions. For instance, in animal embryos, a pH gradient is linked to intestinal stem cell lineage specification and metabolic developmental programming^3,4^. In the plant root epidermis, the extracellular pH (pHe) varies along the longitudinal developmental axis, with the meristematic and differentiation zones distinctively more alkaline than the elongation zone of the root (Fig. 1a)^1,2,5,11^. The plant plasma membrane P-type H^+^-ATPases actively pump H^+^ into the apoplast and are major pHe determinants^9,12^. We analyzed the expression profile of the *Arabidopsis H^+^-ATPases* (*AHAs*) using the Arabidopsis root single cell transcriptome atlas^13^. Among the eleven *AHA* genes present in the *Arabidopsis thaliana* (hereafter Arabidopsis) ecotype Columbia-0 (hereafter Col-0) genome, *AHA2* is the most expressed isoform in the root and is particularly abundant in the trichoblast and atrichoblast cell files that constitute the epidermal cell layer (Fig. 1b, Extended Data Fig. 1a, b). Temporal expression analysis indicated that variation in *AHA2* transcripts accumulation does not correlate with epidermis pHe (Fig. 1c). We thus hypothesize that epidermis pHe is developmentally regulated by additional molecular factors regulating AHAs. We used the *AHA2* temporal expression profile along the longitudinal developmental axis of the atrichoblast and trichoblast cell files to isolate co-regulated genes (Fig 1c). We identified 1985 and 3711 genes co-regulated with *AHA2* (Pearson Coefficient Correlation (PCC) > 0.7) in atrichoblast and trichoblast respectively (Extended Data Fig. 2, Fig. 1d). One thousand and twenty-seven genes were found in both data sets (Fig. 1d), which were enriched in genes encoding plasma membrane-localized proteins (Extended Data Fig. 3). Among them were several genes encoding REMORIN (REM) proteins, including *REM1.1*, *REM1.2*, *REM1.3* and *REM1.4* (Fig. 1e) which belong to the same phylogenetic group^14^; Biermann et al., 2025). REMs are structural elements of the plasma membrane regulating its nano-organization and are best described for their roles in plant-microbe interactions^15–18^. *REM1.2* and *REM1.3* are among the most abundant transcripts and proteins in Arabidopsis (Extended Data Fig 4), suggesting that they play a core cellular function.

**Figure 1.**
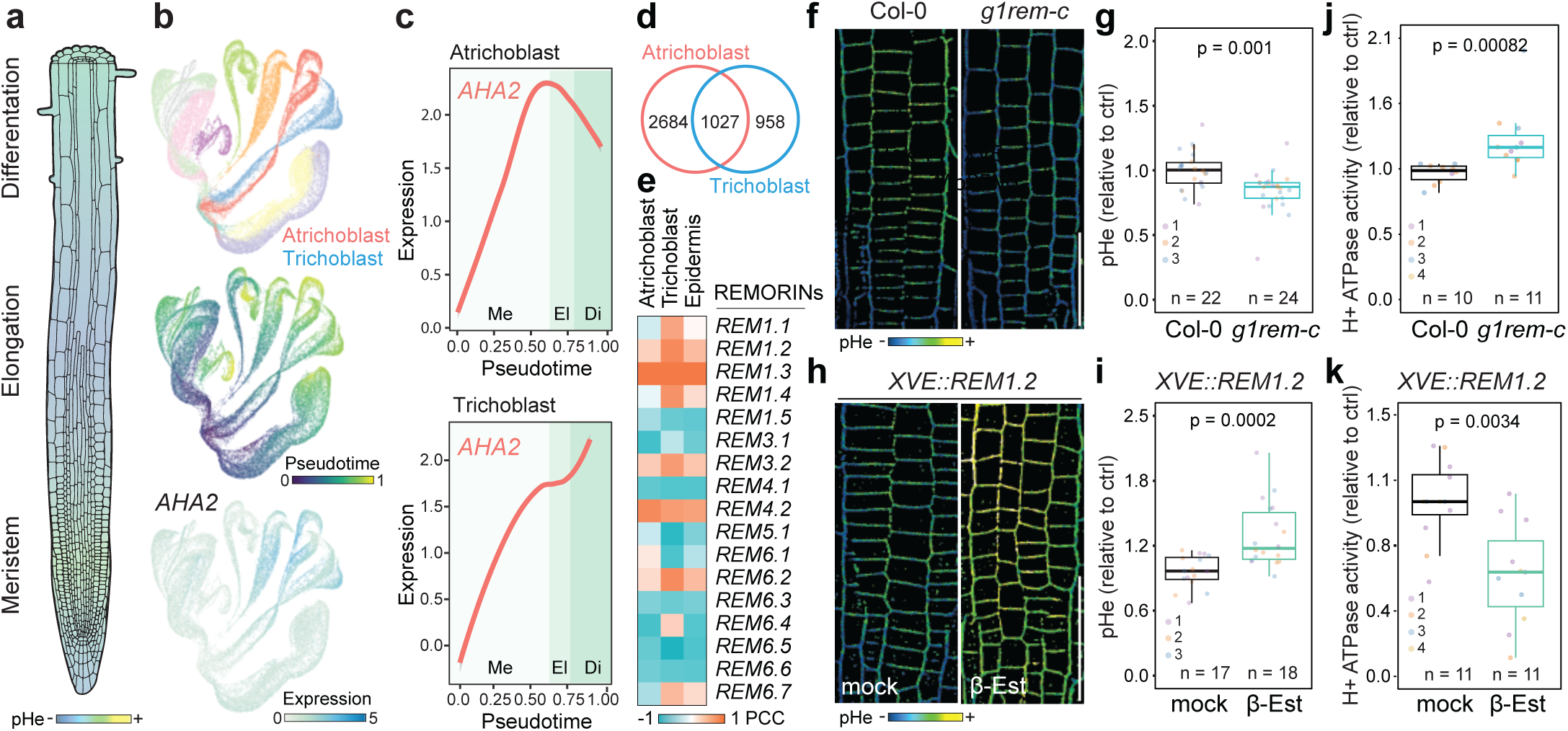
Group 1 *REMORINs* inhibit H^+^-ATPases activity. **a,** Schematic representation of Arabidopsis root tip and of its epidermis pHe. **b,** UMAPs of root tip scRNAseq data ^13^ used in this study. In the UMAPs, cells are labelled according to cell type (top), pseudotime (middle) and *AHA2* expression (bottom). **c,** Pseudotime analysis of *AHA2* expression in the meristem (Me), elongation (El) and differentiation (Di) zones of the atrichoblast and trichoblast cell files. **d,** Venn-diagram of genes co-regulated with *AHA2* in trichoblasts and atrichoblasts. **e**, Pearson correlation coefficient (PCC) of *REMORIN*s expression with *AHA2*. **f** and **h,** Representative confocal microscopy images of HPTS-stained 5-day-old seedling roots (meristem). Scale bars indicate 50 μm. **g** and **i,** Quantification of relative pHe. Data points represent average relative pHe for individual roots (n), colors indicate independent experiments. P values were determined by Mann-Whittney test. **j** and **k**, Quantification of vanadate-sensitive ATP hydrolysis in isolated microsomes of 20-day-old seedlings. Data points represent independent biological samples (n), colors indicate independent experiments. P values were determined by Mann-Whittney test.

To test whether REMs regulate pHe, we generated a quintuple CRISPR mutant (*g1rem-c*) in which the five group 1 Arabidopsis *REMs* genes were edited (Extended Data Fig. 5) and analyzed pHe using the ratiometric pH reporter dye 8-hydroxypyrene-1,3,6-trisulfonic acid trisodium salt (HPTS)^1^. We observed that loss of group 1 *REMs* led to pHe acidification (Fig. 1f, g) indicating that they function as alkalinization factors. Expression of REM1.2 under its native promoter in *g1rem-c* (Extended Data Fig. 5) was sufficient to complement pHe acidification (Extended Data Fig. 6). Therefore, we used REM1.2 as a representative group 1 REM member. We observed that estradiol-inducible expression of REM1.2 (^19^;Extended Data Fig. 5) leads to pHe alkalinization (Fig. 1h, i). To test whether group 1 REMs regulates H^+^-ATPases activity, we measured vanadate-sensitive ATP hydrolysis, a proxy for the activity of plasma membrane-localized H⁺-ATPases^20,21^. We observed an increase in ATP hydrolysis activity in *g1rem-c* mutants (Fig. 1j) suggesting that group 1 REMs are negative regulators of AHAs activity. In good agreement, we observed that REM1.2 overexpression inhibits H^+^-ATPase activity (Fig. 1k). We thus conclude that group 1 REMs promote pHe alkalinization by inhibiting the activity of the plasma membrane H^+^-ATPases.

We observed that GFP-tagged REM1.2 expressed under its native promoter in a *rem1.2* knock-out background^19^ accumulates distinctively in the meristem and differentiation zones of the root, which feature a relative alkaline pHe (Fig. 2a; Extended Data Fig. 7a, b)^2,11^. Live cell imaging of the genetically encoded pHe reporter PM-Apo-Acidin4^11^ alongside with GFP-REM1.2 and GFP-AHA2 confirmed that REM1.2 but not AHA2 protein accumulation correlates with alkaline pHe in the root tip (Fig. 2b, c) – patterns could not be predicted from scRNAseq data (Fig. 1c, Extended Data Fig. 7c-e). In addition, we observed that GFP-REM1.2 accumulates at the base of differentiating trichoblasts and was depleted from the growing tip, variations that correlated with pHe (Extended Data Fig. 8a-c). These observations indicate that REM1.2 marks and defines alkaline pHe zones of the root both at the cellular and tissue level.

**Figure 2.**
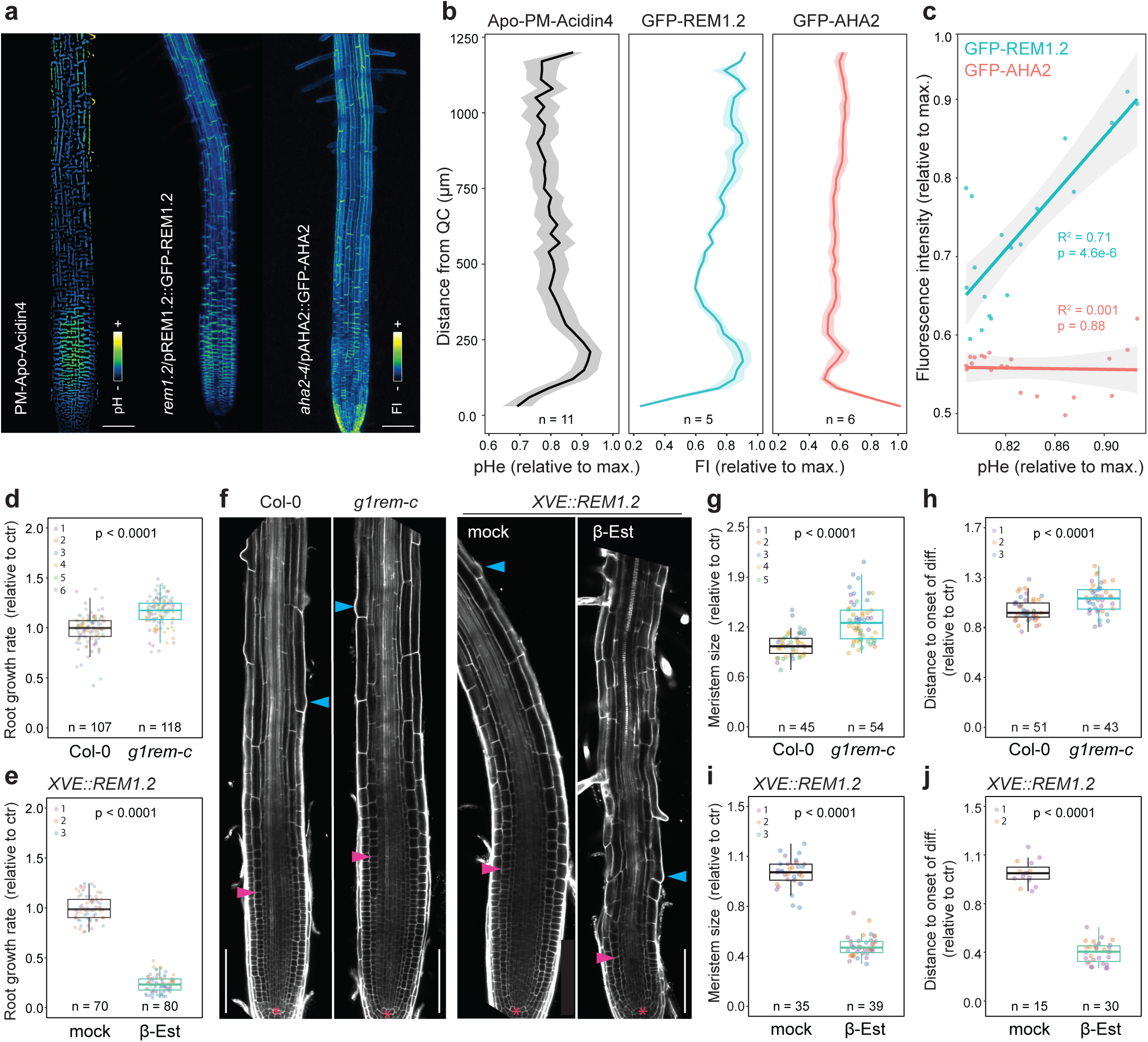
pHe regulates root developmental transitions. **a**, Confocal micrograph z-projections of 5-day-old seedlings root expressing PM-Apo-Acidin4, GFP-REM1.2 or GFP-AHA2. Scale bars indicate 100 μm. **b,** Analysis of the average relative pHe and GFP-REM1.2 or GFP-AHA2 accumulation along root longitudinal growth axis measured on z-projections, (n) indicate the numbers roots measured in three independent experiments. **c**, Linear regression analysis between AHA2-GFP or GFP-REM1.2 accumulation and relative pHe between 100 and 700 µm from the quiescent center. **d** and **e**, Quantification of root growth rate of 5-day-old seedlings, normalized to WT (**d**) and mock treatment (**e**). Data points represent individual roots, colors indicate independent experiments. P values were determined by *t*-test. **f**, Representative root confocal microscopy images of 5-days-old seedlings root stained with propidium iodide. Asterisks indicate the quiescent center, pink and blue arrows mark the ending of the meristem zone and the beginning of the differentiation zone respectively. Scale bars indicate 100 μm. **g** and **i**, Quantification of the meristem size. **h** and **j,** Quantification of the onset of differentiation in 5-day-old seedlings root. Data points represent measurements made in individual roots normalized to WT (**g** and **i**) and mock treatment (**I** and **j**), colors indicate independent experiments. P values were determined by *t*-test.

Genetic and pharmacologic alterations of H^+^-ATPase activity are associated with defects in root development^1,22^. We therefore investigated whether loss and gain of function of group 1 *REMs* alter root development. We observed that *g1rem-c* showed an enhanced root growth rate (Fig. 2d), an increased meristem size, and a delayed differentiation (Fig. 2f-h), which were complemented by the expression of REM1.2 under its native promoter (Extended Data Fig. 6). Conversely, inducing REM1.2 overexpression reduced root growth rate (Fig. 2e), meristem size, and led to a premature differentiation (Fig. 2f, i, j). Altogether, these results show that group 1 *REMs* define root growth rate and developmental transitions. We next questioned whether the regulation of pHe explains the phenotypes of *REMs* gain and loss of function mutants. In agreement, we observed that *g1rem-c* was resistant to the inhibition of meristem size observed in alkaline growth media (pH 7.2) (Extended Data Fig. 9). Further we observed that acidic growth media (pH 4.6) and treatment with fusicoccin (FC), a potent activator of H^+^-ATPases^23^, were sufficient to alleviate the effect of REM1.2 overexpression (Extended Data Fig. 9). Upon H^+^-ATPases inhibition, to compensate for the reduction of the proton motive force required to drive secondary active transport and nutrient uptake, root hair growth increases to enlarge absorbing surface area^24–27^. As predicted by the effect of group 1 REMs on H^+^-ATPase activity (Fig. 1j, k), we observed a decrease in root hair length in *g1rem-c* while REM1.2 overexpression drastically promotes root hair elongation (Extended Data Fig. 8d-g). We conclude that the regulation of pHe by group 1 REMs underlies their role in regulating root development.

We next questioned how the regulation of pHe by REMs regulates root development. Recent evidence showed that pHe is sensed by cell surface peptide-receptor modules and thereby acts as an informative signal in immunity^28,29^ and phloem differentiation^30^. We therefore hypothesize that the modulation of pHe by REMs may regulate cell surface signaling and thus root development. A prominent example is the perception of the growth promoting brassinosteroids (BRs) hormones. Indeed, acidic pH promotes BRs-induced complex formation between the main ligand binding receptor BRASSINOSTEROID INSENSITIVE 1 (BRI1) and its co-receptor BRI1-ASSOCIATED KINASE (BAK1) in vitro^31^. To test if the regulation of apoplast pH by REMs modulates BRI1-BAK1 association in planta, we generated a group 1 *REMs* quintuple CRISPR mutant in a line expressing BRI1-GFP and BAK1-mCherry under the control of their respective promoters (Extended Data Fig. 10). In FRET-FLIM experiments we observed a reduced fluorescence lifetime of BRI1-GFP (Extended Data Fig. 11a, b), indicating an increase in BRI1-GFP/BAK1-mCherry association. BRs regulate root growth in a concentration-dependent manner and excess of BR signaling inhibits meristem size^32,33^. We observed that loss of *group 1 REMs* led to an increased epi-brassinolide (eBL)-induced inhibition of the meristem size (Fig. 3a, b) and of the elongation zone length (Extended Data Fig. 11d), indicative of increased in BR signaling. In good agreement, we observed an increased dephosphorylation of the BR-responsive transcription factor BES1^34^ in *g1rem-c* (Extended Data Figure 11c). We next monitored transcript accumulation of two marker genes negatively regulated by BRs, namely *CPD* and *DWF4*^35^. We observed a decrease in transcript accumulation of both *CPD* and *DWF4* (Extended Data Fig. 11e). Conversely, we observed that *REM1.2* overexpression led to hyposensitivity to exogenous eBL treatment (Fig. 3c, d; Extended Data Fig. 11f) and increased *CPD* transcript accumulation (Extended Data Fig. 11g). We conclude that group 1 REMs inhibit BR signaling, as predicted by their effect on pHe.

**Figure 3.**
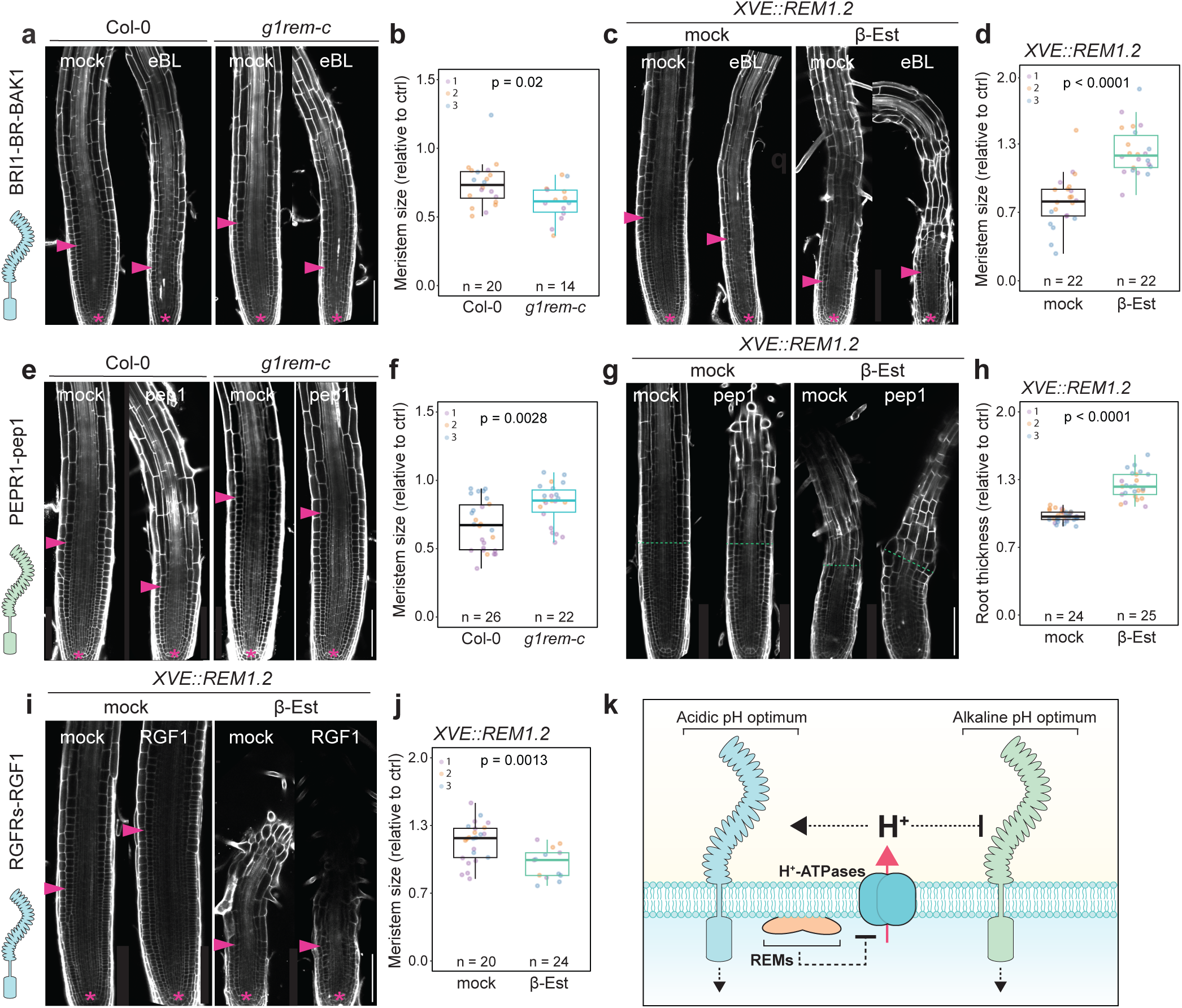
*REMORINs* modulate cell surface ligands responsiveness. Confocal micrographs of roots of 5-day-old seedling germinated on media containing 1 nM eBL (**a**) or 5 nM eBL (**c**) or treated with 10 nM pep1 for 12 h (**e** and **g**) or 20 nM RGF1 for 24 h (**i**) or corresponding control mock solutions and stained propidium iodide. Scale bars indicate 100 µm. Quantification of meristem size or root thickness of roots of 5-day-old seedling germinated on media containing 1 nM eBL (**b**), or 5 nM eBL (**d**), or treated with 10 nM pep1 for 12 h (**f and h**) or 20 nM RGF1 for 24 h (**j**) or corresponding control mock solutions. Data points represent measurements made in individual roots normalized to the mock treatments (without exogenous ligand treatment) in WT, *g1rem-c*, *XVE::REM1.2* (mock) and *XVE::REM1.2* (β-Est) conditions, colors indicate independent experiments. P values were determined by Mann-Whitney test. **k**, schematic proposed representation of the effect of pHe regulation by group 1 *REMs* on cell surface signaling, see also Extended Data Fig. 11.

In contrast to the BRI1-BR-BAK1 complex, binding of the damage-associated molecular pattern plant elicitor peptide 1 (pep1) to its receptor PLANT ELICITOR PEPTIDE RECEPTOR 1 (PEPR1) and consequently pep1-induced signalling is enhanced at relative alkaline pH^29,36^. We observed that the inhibition of meristem size by pep1^29^ was reduced in *g1rem-c* (Fig. 3e, f), while pep1-induced root swelling^37^ was enhanced upon REM1.2 overexpression (Fig. 3g, h). Furthermore, we observed that the inhibition of pep1-induced MAPK phosphorylation by acidic pH media was suppressed by REM1.2 overexpression (Extended Data Fig. 11h). *REM1.2* overexpression also inhibited responsiveness to exogenous ROOT GROWTH FACTOR 1 (RGF1^38^) treatment (Fig. 3i, j) in agreement with the inhibition of its perception by RGFR1 under pronounced alkaline conditions^29^. In contrast, loss of group 1 *REMs* did not affect RGF1 responsiveness (Extended Data Fig. 11i, j). In addition, REM knock-out and over-expression lines showed altered responsiveness to the root RAPID ALKALINIZATION FACTOR 1 (RALF1)^39^ and RALF22^40^ (Extended Data Fig. 11k) consistent with RALF binding’s pH optimum to their cell wall-located receptors^41^. Altogether, these observations indicate that the regulation of pHe by REMs modulates various cell surface signaling pathways. Furthermore, we observed that loss and gain of function REM mutants were affected in cell wall pectin esterification status and that REM1.2 overexpression inhibits bulk endocytic fluxes (Extended Data Fig. 12). These observations resonate with the effect of pH on the inhibition of PECTIN METHYLESTERASE (PME) by PME INHIBITOR in vitro^42^ and on endocytosis in animal cells^43,44^. We conclude that regulation of pHe by REMs modulate various cell surface processes underscoring their role in development.

We next performed a genome-wide analysis of REMs occurrence in the green lineage. We found that REMs emerged within Streptophyte algae, with REM members encoded in the genome of the Charophyceae *Chara braunii* and conserved in all land plants (Fig. 4a, Extended Data Fig. 13). Interestingly, we observed that the occurrence of *REM* genes correlates with complex plant body structure. Indeed, REMs are absent in the Coleochaetophyceae and Zygnematophyceae Streptophyte algae investigated (Extended Data Fig. 13). Considering molecular phylogenetic studies^45–47^, these observations suggest that REMs correspond to key evolutionary innovations present in the land plants common ancestor. We next asked whether the inhibition of H^+^-ATPases by REMs is evolutionary conserved. For this purpose, we generated knockout and overexpression lines of Mp*REM1* the closest orthologue of *Arabidopsis* group 1 REMs in *Marchantia polymorpha* (hereafter Marchantia) (Fig. 4b; Extended Data Fig. 14)(Biermann et al., 2025). Marchantia belongs to the bryophytes, the sister lineage to vascular plants lacking a true vasculature and root system^48^. We observed that loss of Mp*REM1* led to pHe acidification and an increase in H^+^-ATPases activity (Fig. 4c-e). Conversely, we observed that Mp*REM1* overexpression led to pHe alkalinization and to a decrease in PM H^+^-ATPases activity (Fig. 4f-h). These observations show that the inhibition of H^+^-ATPases by REMs is conserved in Marchantia, deeply conserved in the REM protein family, and predates the emergence of the root system itself.

**Figure 4.**
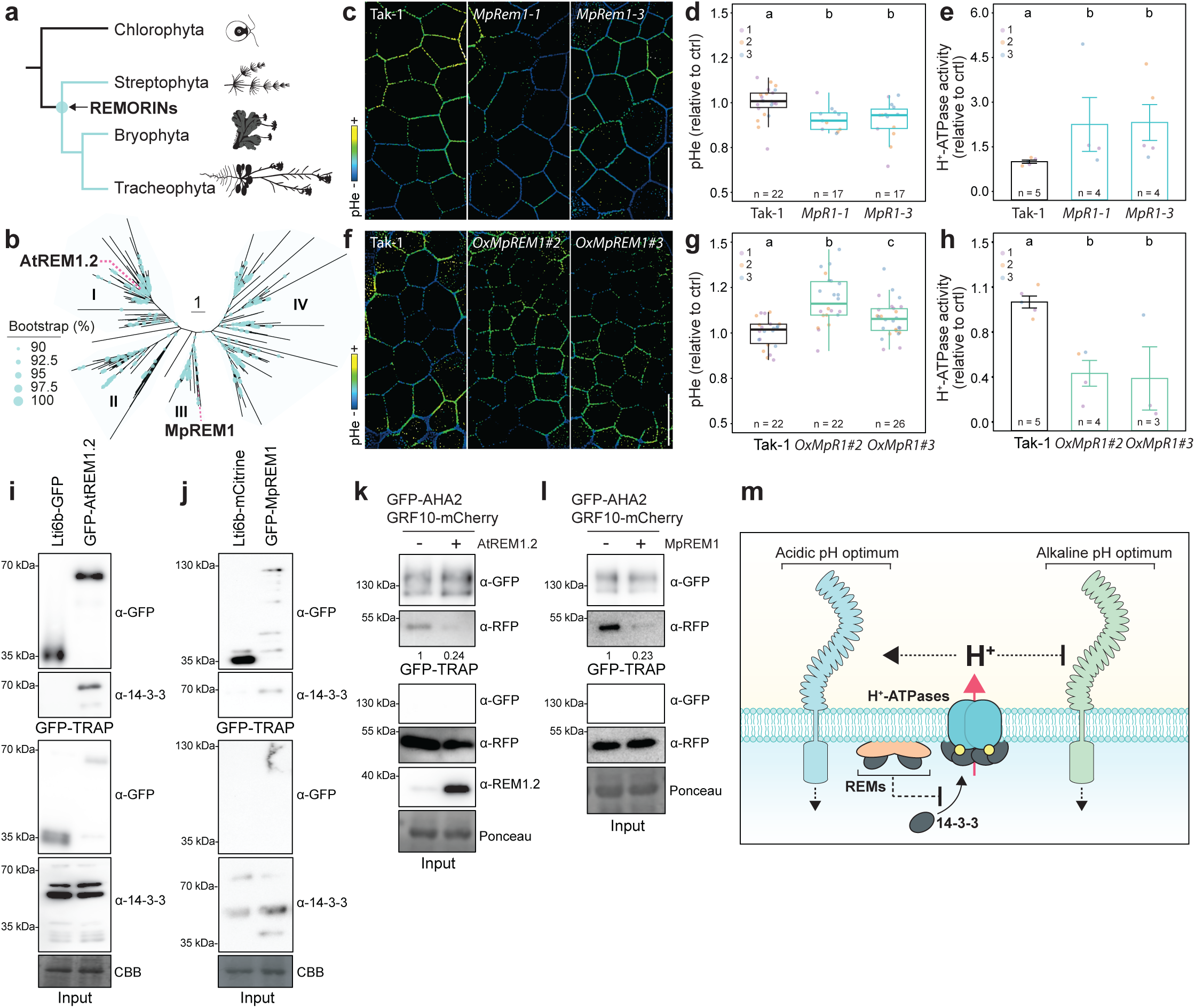
REMORINs are conserved evolutionary innovation. **a,** Schematic representation of the emergence of REMs in the green lineage, see also Extended Data Figure 13. **b**, Maximum likelihood tree of full-length REM proteins from Biermann et al. (2025), likelihood equal −377021.439, branch support is indicated in % from 2000 bootstraps, scale bar indicates substitution per site. The position of AtREM1.2 and MpREM1 is indicated. **c** and **f**, Representative confocal micrographs of 1-day-old gemmalings thallus epidermis stained with HPTS. Scale bars indicate 50 μm. **d** and **g,** Quantification of relative pHe, Data points represent measurement from (n) individual gemmalings, normalized to the control Tak-1, colors indicate independent experiments. Conditions which do not share a letter are statistically different in Dunn’s multiple comparison test (p < 0.05). **d**, Quantification of vanadate-sensitive ATP hydrolysis in isolated microsomes of 20-day-old Marchantia thallus, normalized to WT Tak-1. Data points represent measurements from (n) individual biological replicates, each corresponding to 5-4 pooled plants, normalized to the control Tak-1, colors indicate independent experiments. Conditions which do not share a letter are statistically different in Dunn’s multiple comparison test (p < 0.05). **h,** Immunoprecipitation experiment using Col-0/p35S::Lti6b-GFP and *rem1.2*/pREM1.2::GFP-REM1.2 Arabidopsis seedlings. Membranes were probed with anti-GFP and anti-14-3-3 antibodies. CBB staining is shown to assess protein loading. Similar observations were made in at least three independent experiments. **i,** Immunoprecipitation experiments using Tak-1/p35S::LTI6B-mCitrine and MpRem1-1/p35S::GFP-MpREM1 *Marchantia* plants. Membranes were probed with anti-GFP and anti-14-3-3 antibodies. CBB staining is shown to assess protein loading. Similar observations were made in at least three independent experiments. **k** and **l,** Immunoprecipitation experiments of pUb10::GFP-AHA2 transiently co-expressed with pUb10::GRF10-mCherry and pUb10::AtREM1.2 (**k**) or pUb10::MpREM1 and treated with 1 μM FC for 1 h (**l**) in *Nicotiana benthamiana* leaves. Membranes were probed with anti-GFP, anti-RFP and anti-REM1.2 antibodies. Numbers indicate chemiluminescence intensity of co-immunoprecipitated GRF10-mCherry relative to the condition – AtREM1.2 or – MpREM1. *MpREM1* expression has been controlled by RT-qPCR (Extended Data Fig. 14c). Ponceau staining is shown to assess protein loading. Similar observations were made in at least three independent experiments. **k**, Working model for the regulation of H^+^-ATPases activity and cell surface signaling by REMs.

We next investigated how REMs regulate the activity of H^+^-ATPases. In plants, H^+^-ATPases are commonly regulated by phosphorylation of their inhibitory C-terminal regulatory domain, notably on the conserved penultimate residue (Threonine 947 in AHA2) allowing for binding of 14-3-3 dimers which favor the release of the inhibitory domain thus promoting H^+^-ATPases activation^24,49^. AtREM1.2 was previously reported to associate with 14-3-3 dimers and in particular the 14-3-3 GENERAL REGULATORY FACTOR 10 (GRF10)^19^. In good agreement, in co-immunoprecipitation experiments, we observed that GFP-AtREM1.2 associates with Arabidopsis and *Nicotiana benthamiana* endogenous 14-3-3 (Fig. 4i, Extended Data Fig. 15a). Further, we observed that GFP-MpREM1 co-immunoprecipitated with a 14-3-3 homolog in Marchantia (Fig. 4j), indicating that the association between REMs and 14-3-3 protein is evolutionary conserved. Further, both AtREM1.2 and MpREM1 were co-immunoprecipitated with AHA2, suggesting that REMs may associate with 14-3-3 proteins in the vicinity of H^+^-ATPases (Extended Data Fig. 15b, c). We hypothesized that REMs inhibit 14-3-3/H^+^-ATPase association. In good agreement, we observed that AtREM1.2 overexpression in *Nicotiana benthamiana* inhibited the association between GFP-AHA2 and GRF10-mCherry (Fig. 4k). Similarly, we observed that MpREM1 overexpression led to a decrease in AHA2-GRF10 association (Fig. 4l, Extended data Fig. 15b). Altogether, these observations indicate that REMs bind to 14-3-3 proteins to inhibit H^+^-ATPases, providing a mechanism for the conserved role of REMs in regulating H^+^-ATPases. In sum, this study demonstrates that REM proteins act as evolutionarily conserved inhibitors of plasma membrane H⁺-ATPases, thus identifying an elusive regulator of pHe and defining its role in development. Notably, Arabidopsis group 1 REMs modulate various cell surface processes in accordance with their pH sensitivity (Fig. 3). These observations support that pHe acts as an informative signal^28–30^ whose influence extends beyond the classical acid growth theory^50,51^. We envision that the inhibition H^+^-ATPase by REMs provides the means to define spatial pHe patterns that supported the increase in complexity of plant body architecture over evolutionary time. Further work is needed to understand how the REM/14-3-3/H^+^-ATPase module is integrated into stress and development signaling and to clarify its role(s) during plant evolution.

## Acknowledgement

We thank all members of the NanoSignaling Laboratory for fruitful discussions and comments on the manuscript. We thank Emmanuelle Bayer, Steven Huber, Yansong Miao, Xu Chen, Jiří Friml and Nadine Paris for providing materials. We thank Elke Barbez for sharing detailed protocols for HPTS staining and imaging. We thank Nga Pham for technical assistance. Confocal microscopy was performed at the ZMBP microscopy facility of the University of Tübingen. This research was supported by the Deutsche Forschungsgemeinschaft (DFG) grants A08-SFB1101 and B01-TRR356 to J.G., the European Molecular Biology Organization (postdoctoral fellowship EMBO LTF 438-2018, to J.G.), the European Research Council under the grant agreement 773153 (immuno-PEPTALK to C.Z.) and by the Technical University of Munich.

## Author contributions

Conceptualization and project administration, J.G., funding acquisition, C.Z., J.G.; investigation, K.XH., M.v.A., D.B., A.P., P.P.; supervision, J.G.; resources, F.K., N.P., N.F., I.M., C.Z.; writing – original draft, K.XH, J.G.; writing – review and editing, D.B., M.v.A., I.M., F.K., C.Z., C.O., M.T., K.XH, J.G.

---

**Extended Data Figure 1.**
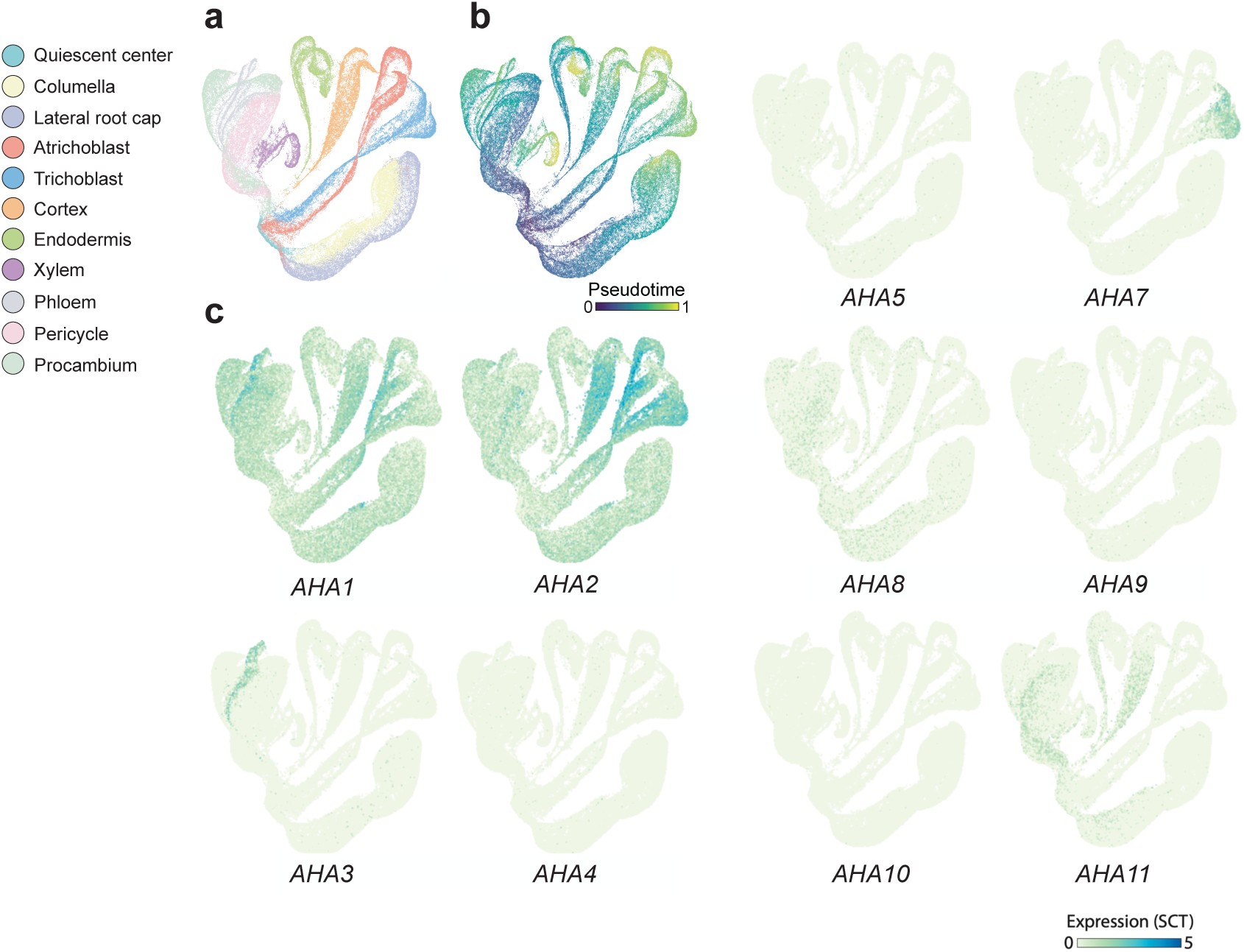
Analysis of *AHAs* expression in Arabidopsis ScRNAseq atlas. UMAPs of root tip ScRNAseq data^13^ labelled according to cell type (a), pseudotime (b) and *AHAs* expression (c). Note that *AHA6* transcripts were not detected.

**Extended Data Figure 2.**
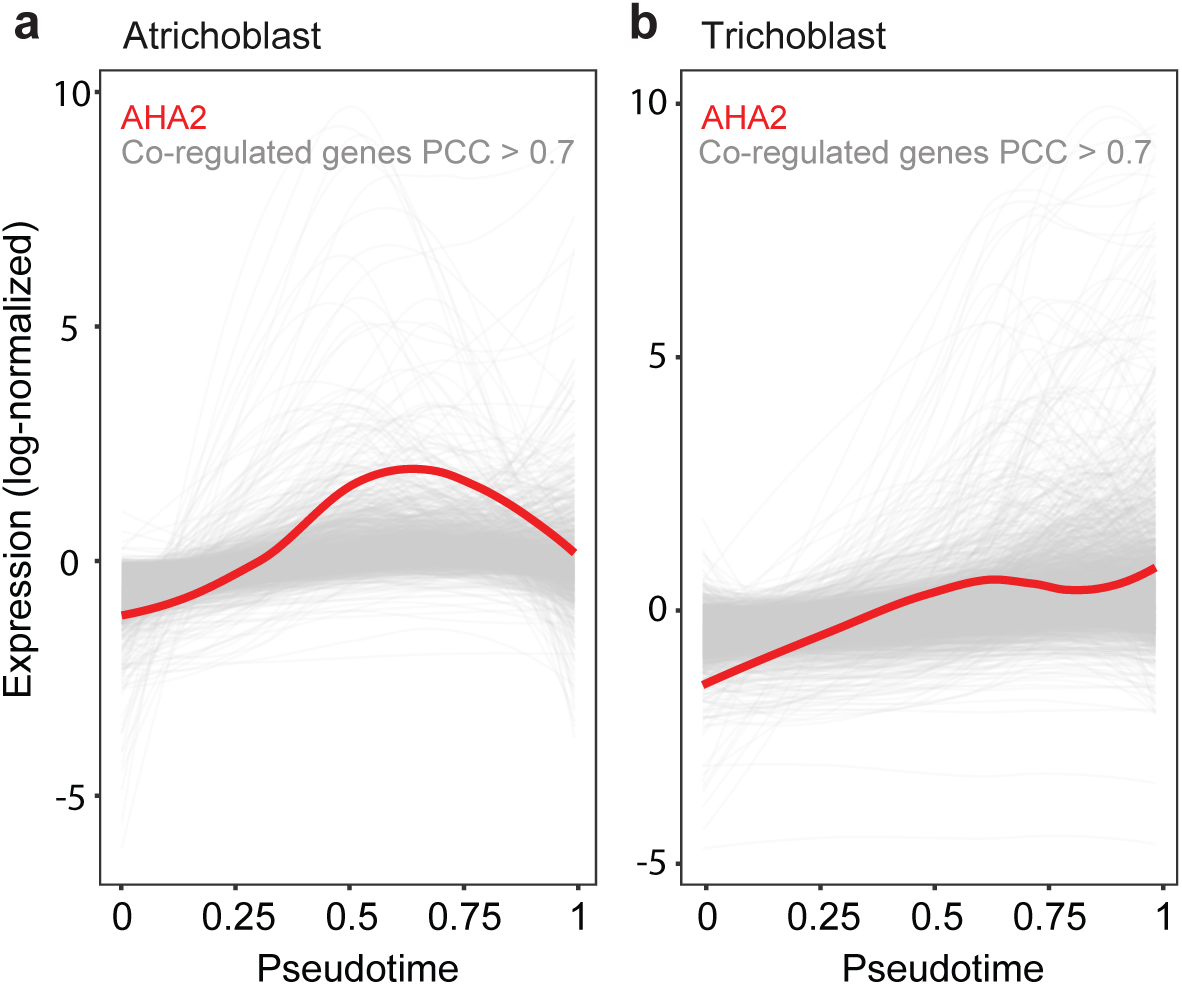
Identification of AHA2 co-regulated genes. Graphs represent temporal expression of *AHA2* and co-regulated transcripts (PCC > 0.7) in atrichoblast (a) and trichoblast (b) cell files.

**Extended Data Figure 3.**
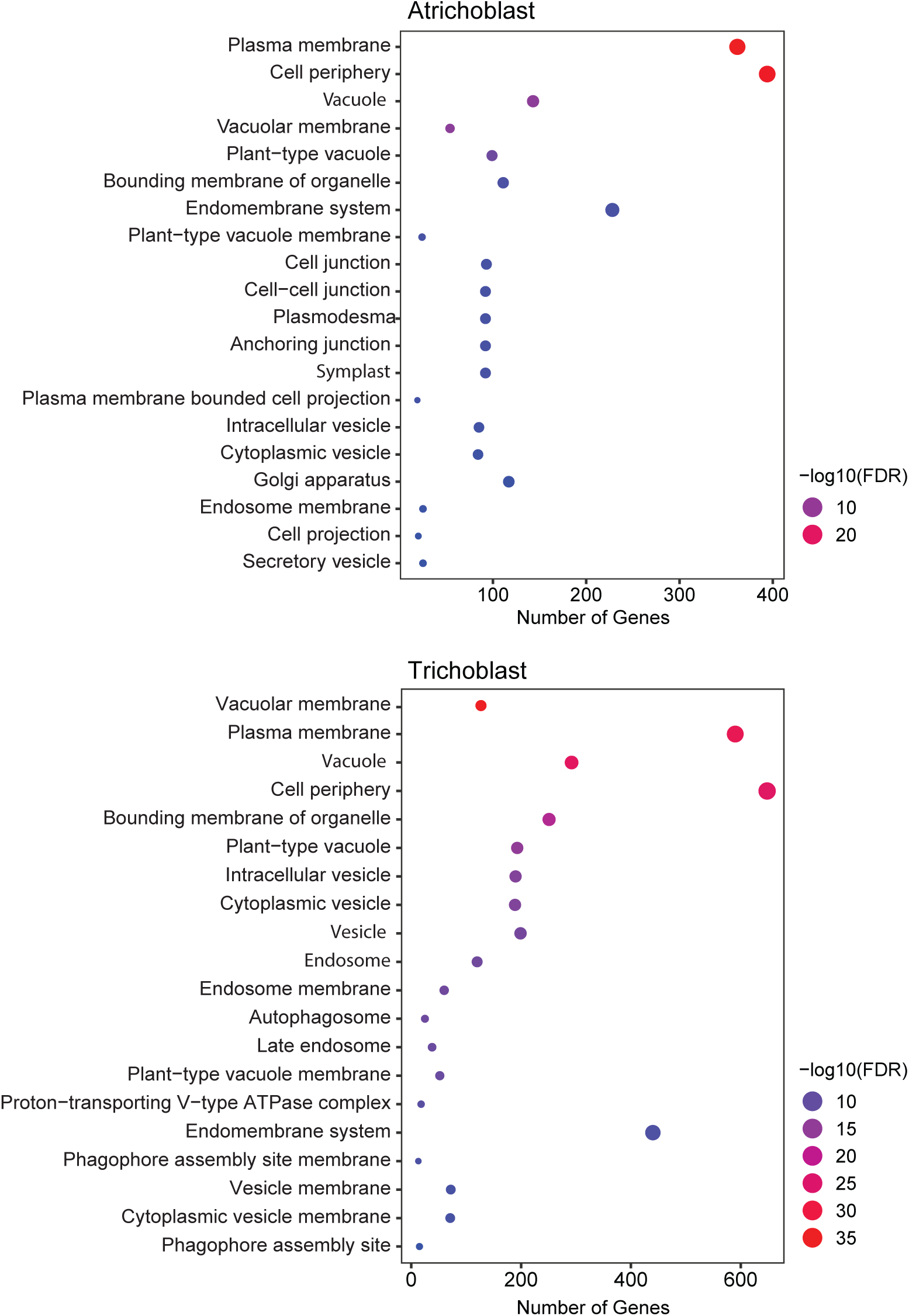
Ontology of *AHA2* co-regulated genes. Gene ontology enrichment analyses of the cellular compartments represented by the proteins encoded by the *AHA2* co-regulated genes in atrichoblast (top) and trichoblast cells.

**Extended Data Figure 4.**
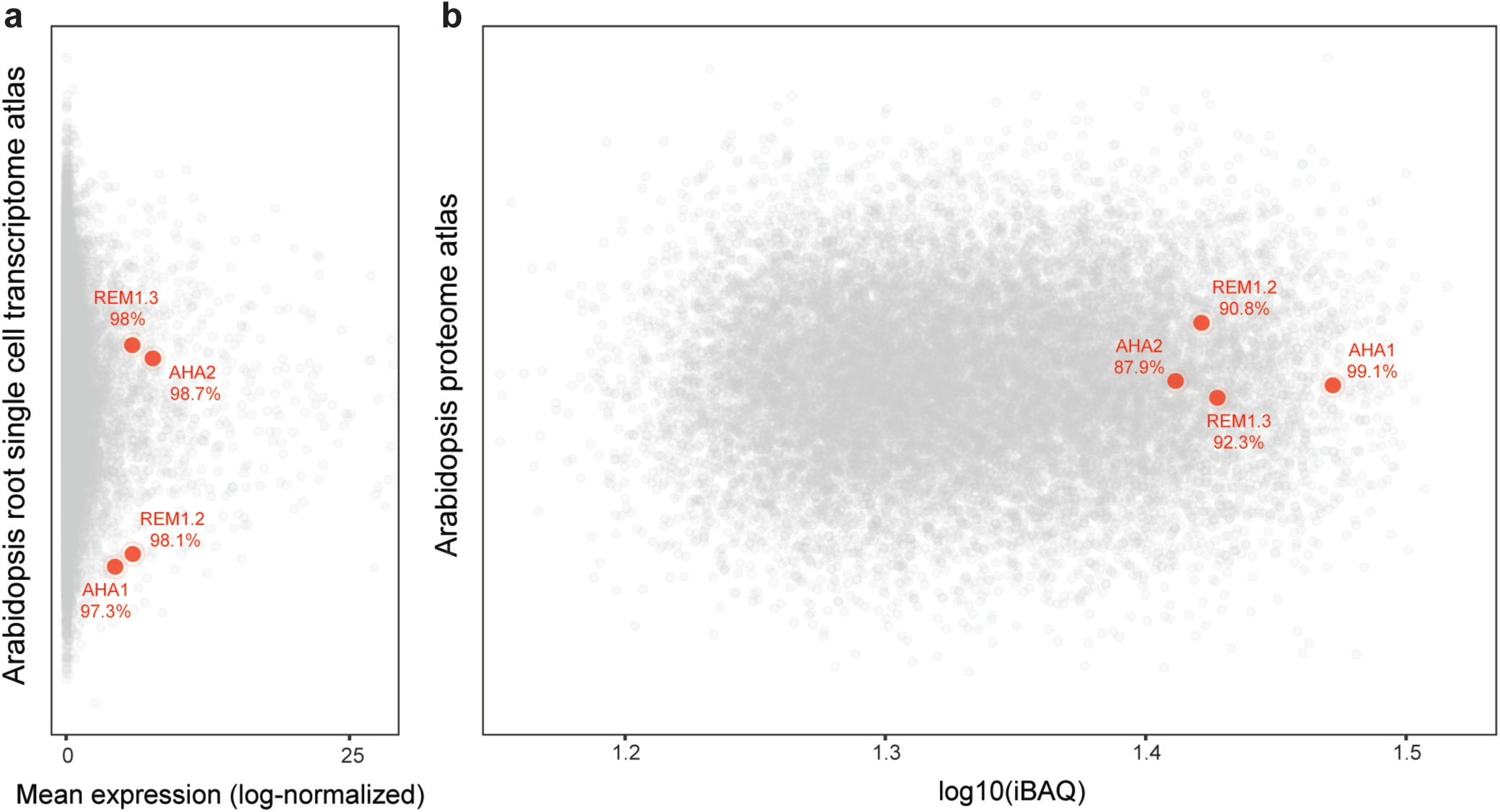
Analysis of REM1.2 and REM1.3 transcript and protein abundance. **a**, Arabidopsis root single cell transcriptome atlas^13^. **b**, Intensity-based absolute quantification (iBAQ) of Arabidopsis proteins^52^. The position of REM1.2, REM1.3, AHA1 and AHA2 transcripts and proteins is indicated in red, the percentages indicate the percentile scores of the corresponding transcript/protein, e.g., *REM1.2* transcripts are among the top two-percent most abundant transcripts in Arabidopsis root single cell transcriptome atlas.

**Extended Data Figure 5.**
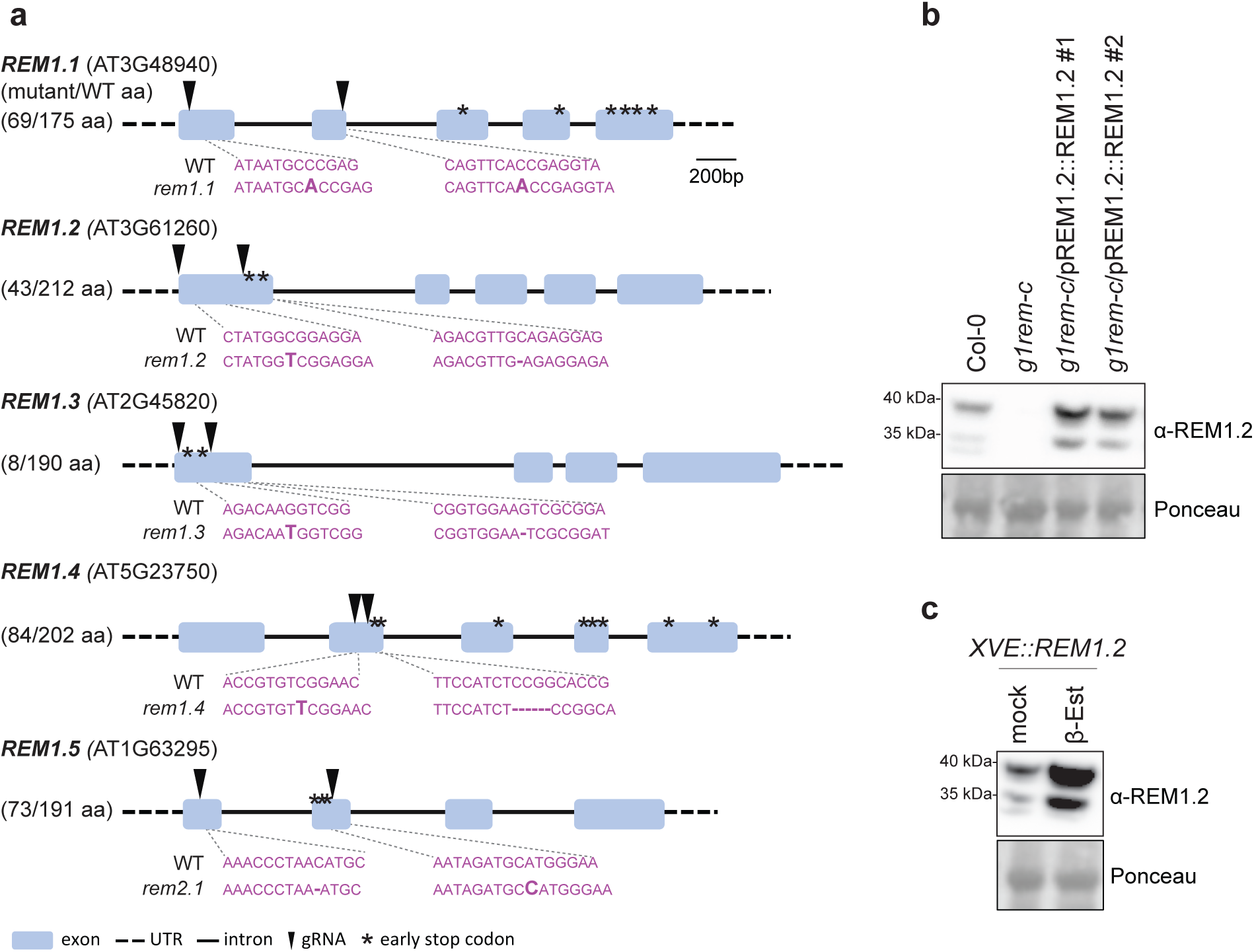
Analysis of group 1 *REMs* genetic material. **a**, Representation of group 1 *REMORIN* genes loci. Black arrows indicate the position targeted by gRNAs. Sequences of *REM* genes in WT and *g1rem-c* are shown in pink, editing events are shown in bold, asterisk indicates early stop codons caused by editing events. Amino acid (aa) numbers indicate the protein and protein fragment expected to be produced WT and *g1rem-c* plants. **b,** Western blot analysis of REM1.2 protein accumulation in WT, *g1rem-c*, and *g1rem-c* complementation lines. The membrane was probed with anti-REM1.2 and stained with Ponceau to assess protein loading. **c**. Western blot analysis of REM1.2 accumulation upon inducible expression upon treatment with 0.5 μM β-Estradiol or corresponding mock control (EtOH). The membrane was probed with anti-REM1.2 and stained with Ponceau to assess protein loading.

**Extended Data Figure 6.**
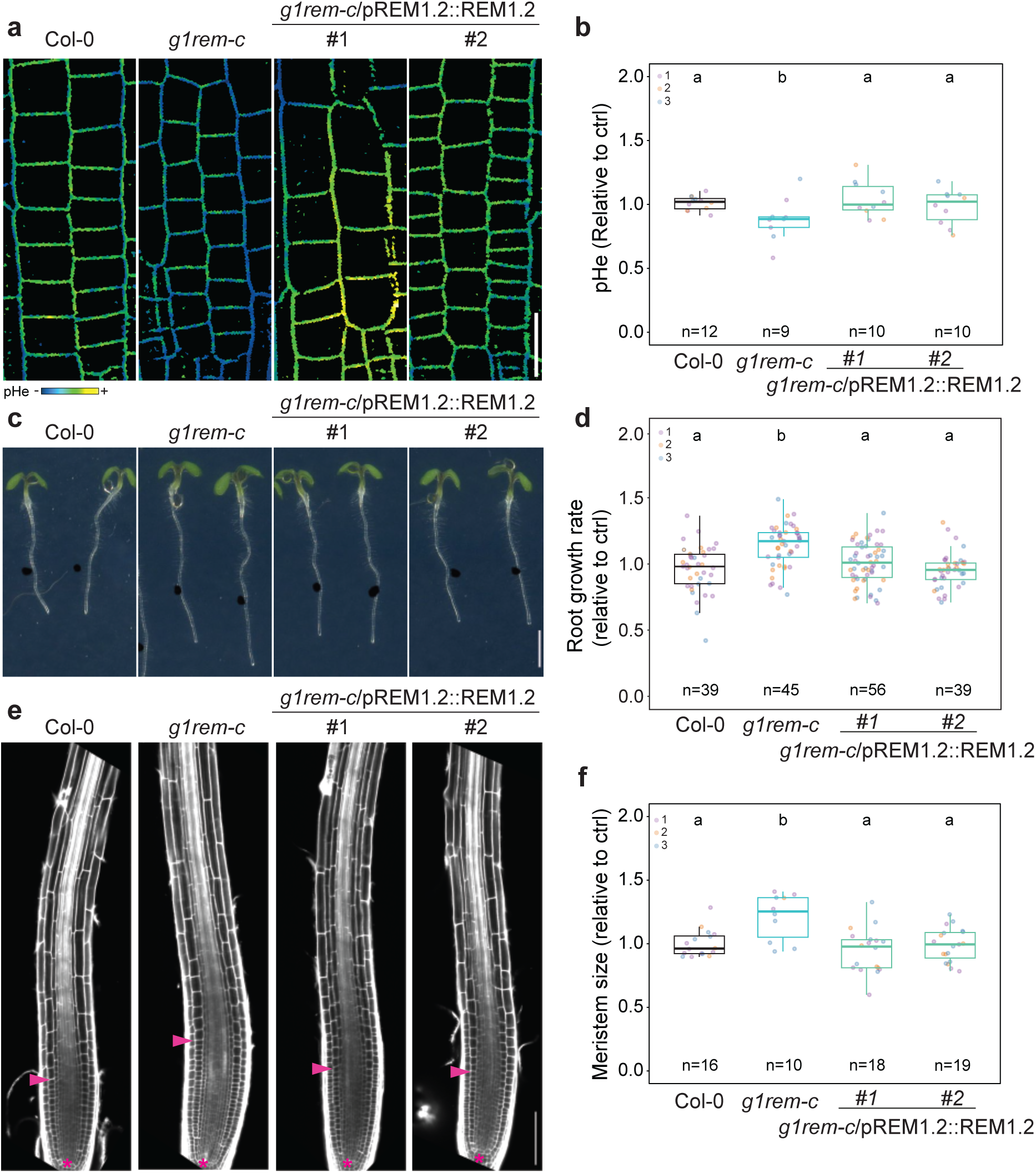
Phenotypic analysis of *g1rem-c* complementation lines. **a,** Representative confocal microscopy images of HPTS-stained 5-day-old seedling roots (meristem cells). Scale bar indicates 20 μm. **b,** Quantification of relative pHe. Data points represent average relative pHe for individual roots (n), colors indicate independent experiments. Conditions which do not share a letter are statistically different in Dunn’s multiple comparison test (p < 0.05). **c**, Representative images of 5-day-old seedlings grown on ½ MS medium. The scale bar indicates 0.2 cm. **d,** Quantification of root growth rate, data points are measurement of individual roots (n) normalized to WT, colors indicate independent experiments. Conditions which do not share a letter are statistically different in Kruskal-Wallis test (p < 0.05). **e,** Representative confocal microscopy images of 5-day-old seedling roots stained with propidium iodide. Asterisks indicate the quiescent center and arrows mark the end of the meristematic zone. Scale bar indicates 100 μm. **f**, Quantification of the meristem size for individual roots (n) normalized to WT, colors indicate independent experiments. Conditions which do not share a letter are statistically different in Dunn’s multiple comparison test (p < 0.05).

**Extended Data Figure 7.**
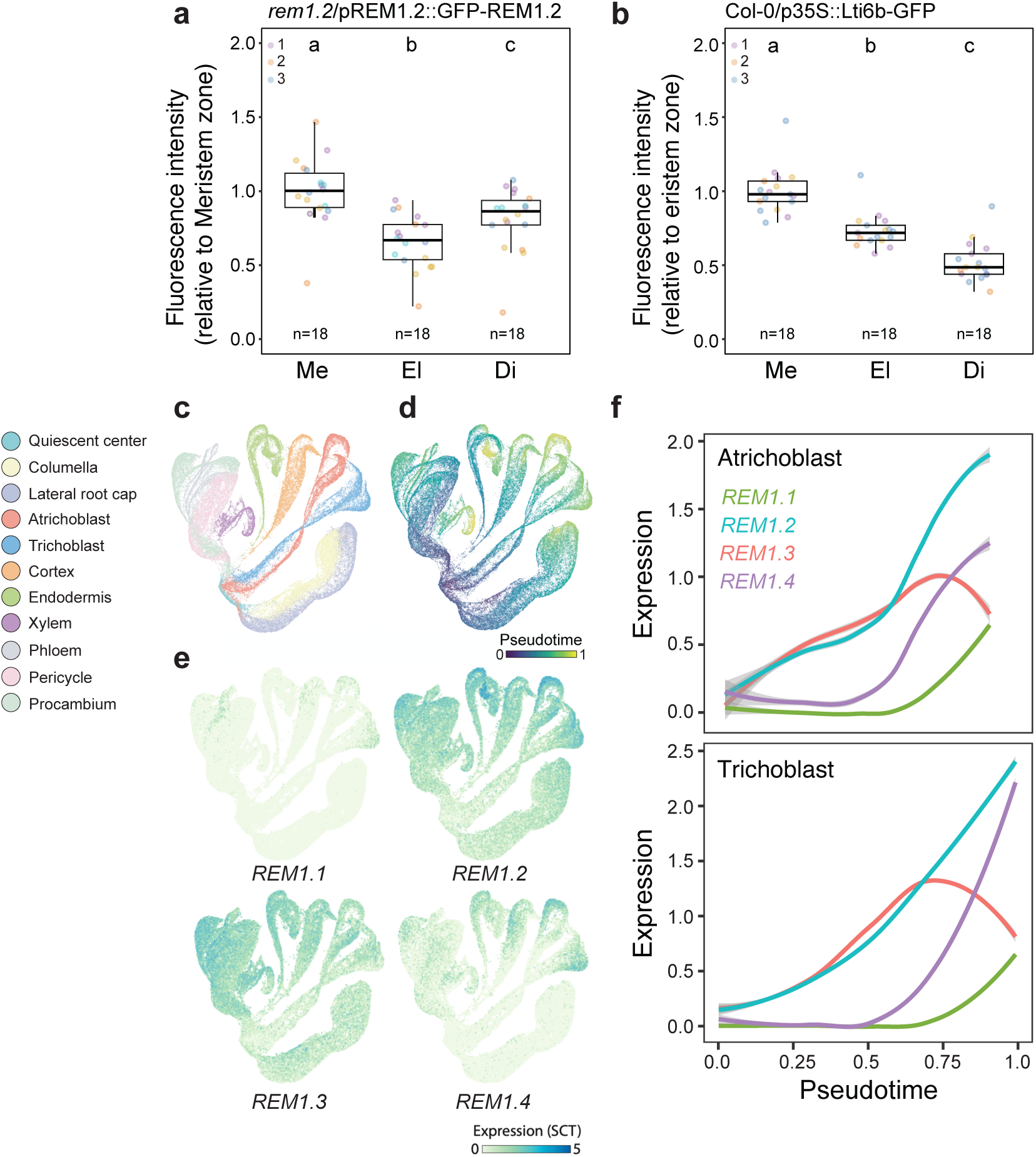
Analysis of *REMs* expression in Arabidopsis ScRNAseq atlas. **a**-**b**, Quantification of the relative fluorescence intensity of GFP-REM1.2 (**a**), and Lti6b-GFP (**b**) in root tip of 5-day-old seedlings, data points are measurement of individual roots (n) normalized to WT, colors indicate independent experiments. Conditions which do not share a letter are statistically different in Kruskal-Wallis test (p < 0.05). Meristem (Me), elongation zone (El) and differentiation zone (Di). **c**-**e**, UMAPs of root tip ScRNAseq data^13^ labelled according to cell type (**c**), pseudotime (**d**) and *REMs* expression (**e**). **f**, Pseudotime analysis of REMs expression in atrichoblast (top) and trichoblast (bottom) cell files.

**Extended Data Figure 8.**
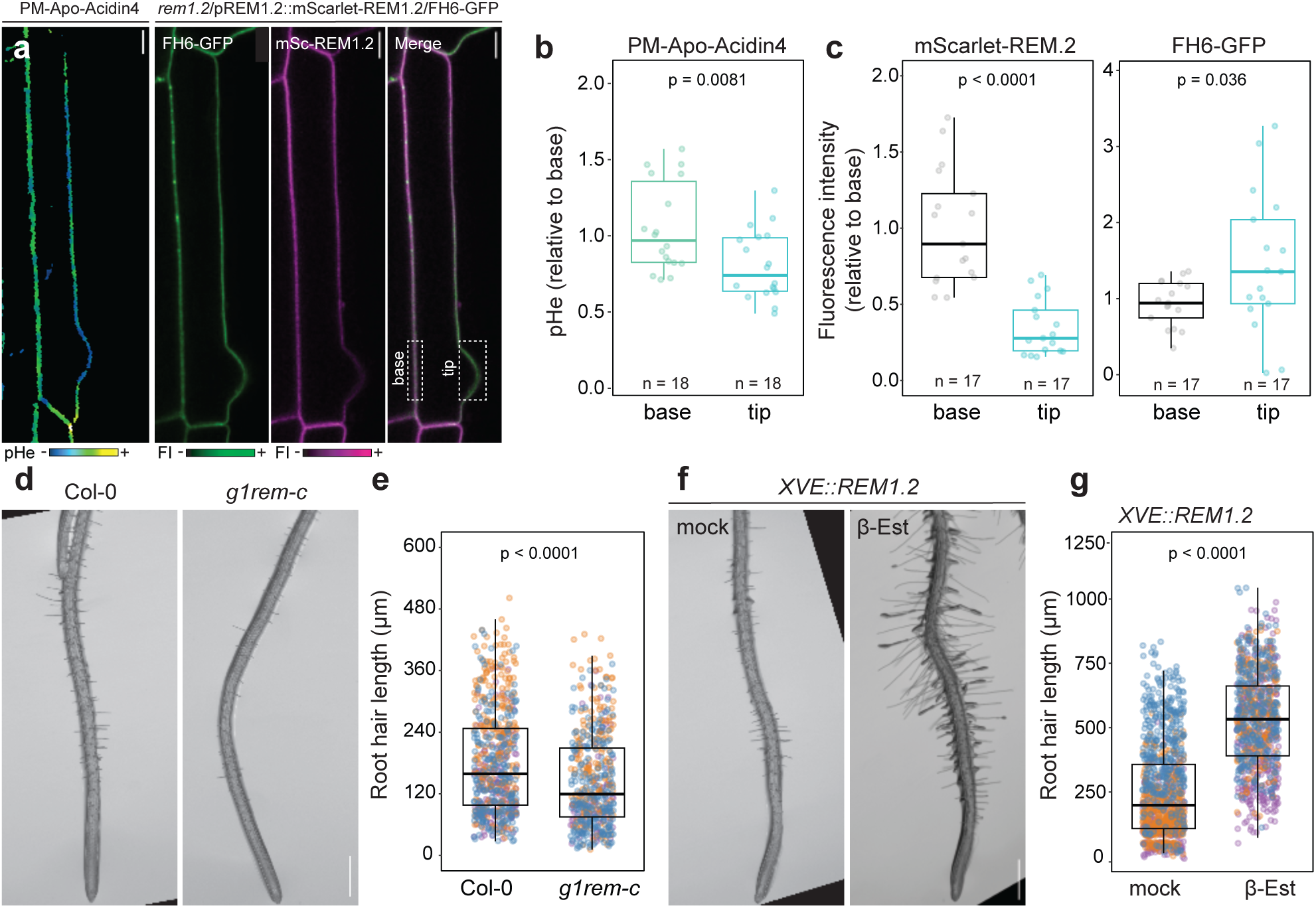
Analysis of group 1 *REMs* in Arabidopsis root hairs. **a,** Representative confocal microscopy images of trichoblast cells presenting an emerging root hair in Col-0/Apo-PM-Acidin4 and *rem1-2*/pREM1.2::mScarlet-AtREM1.2/AtFH6-GFP stable transgenic lines. Scale bar indicates 10 μm. **b,** Quantifications of relative pHe at the base and tip of emerging root hairs, n indicates the number of trichoblast cells analysed. P value reports Mann-Whitney test. **c,** Quantification of the relative accumulation of mScarlet-REM1.2 and FH6-GFP at the tip and base of emerging root hairs, n indicates the number of trichoblast cells analysed. P value reports Mann-Whitney tests. **d** and **f,** Stereoscopic images of 5-day-old seedling roots. Scale bar indicates 1 mm. **e** and **g,** Quantification of root hair length, data points are individual root hair measurement, colors indicate independent experiments. P values report *t*-tests.

**Extended Data Figure 9.**
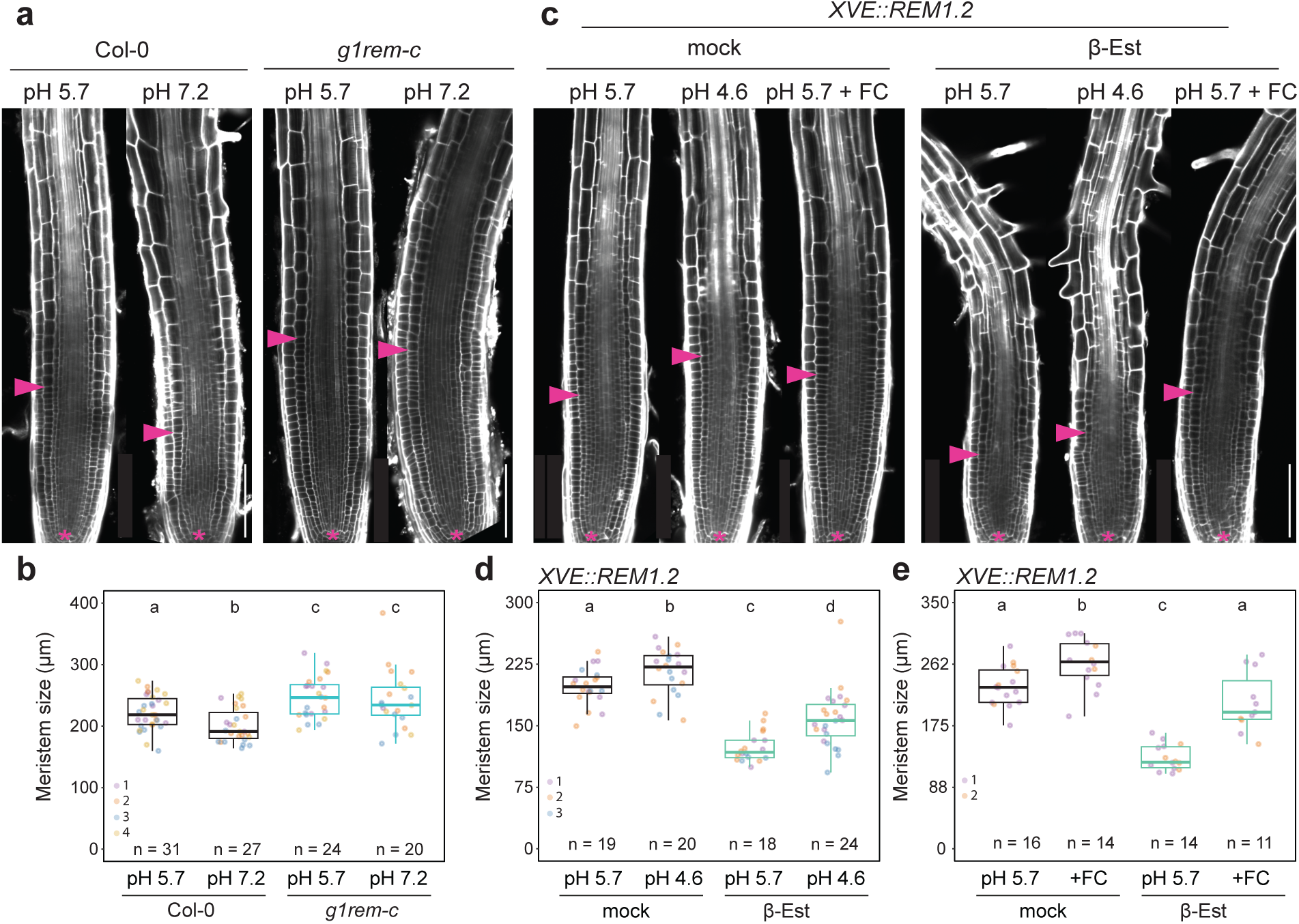
Analysis of group 1 *REMs* gain and loss of function mutants in varying pH conditions. **a** and **c**, Representative confocal microscopy images of 5-day-old seedling roots incubated for 12 h in liquid ½ MS media adjusted to pH 4.6, pH 5.4, pH 7.2 and/or treated with fusicoccin (FC) or the corresponding mock control and stained with propidium iodide. Asterisks indicate quiescent center, arrows mark the end of the meristem. Scale bar indicates 100 μm. **b, d** and **e**, Quantification of the meristem size of 5-day-old seedling roots treated for 12 h in liquid ½ MS with the indicated conditions. Data points how measurements of individual (n) roots, colors indicated independent experiments. Conditions which do not share a letter are statistically different in Kruskal-Wallis test (p < 0.05).

**Extended Data Figure 10.**
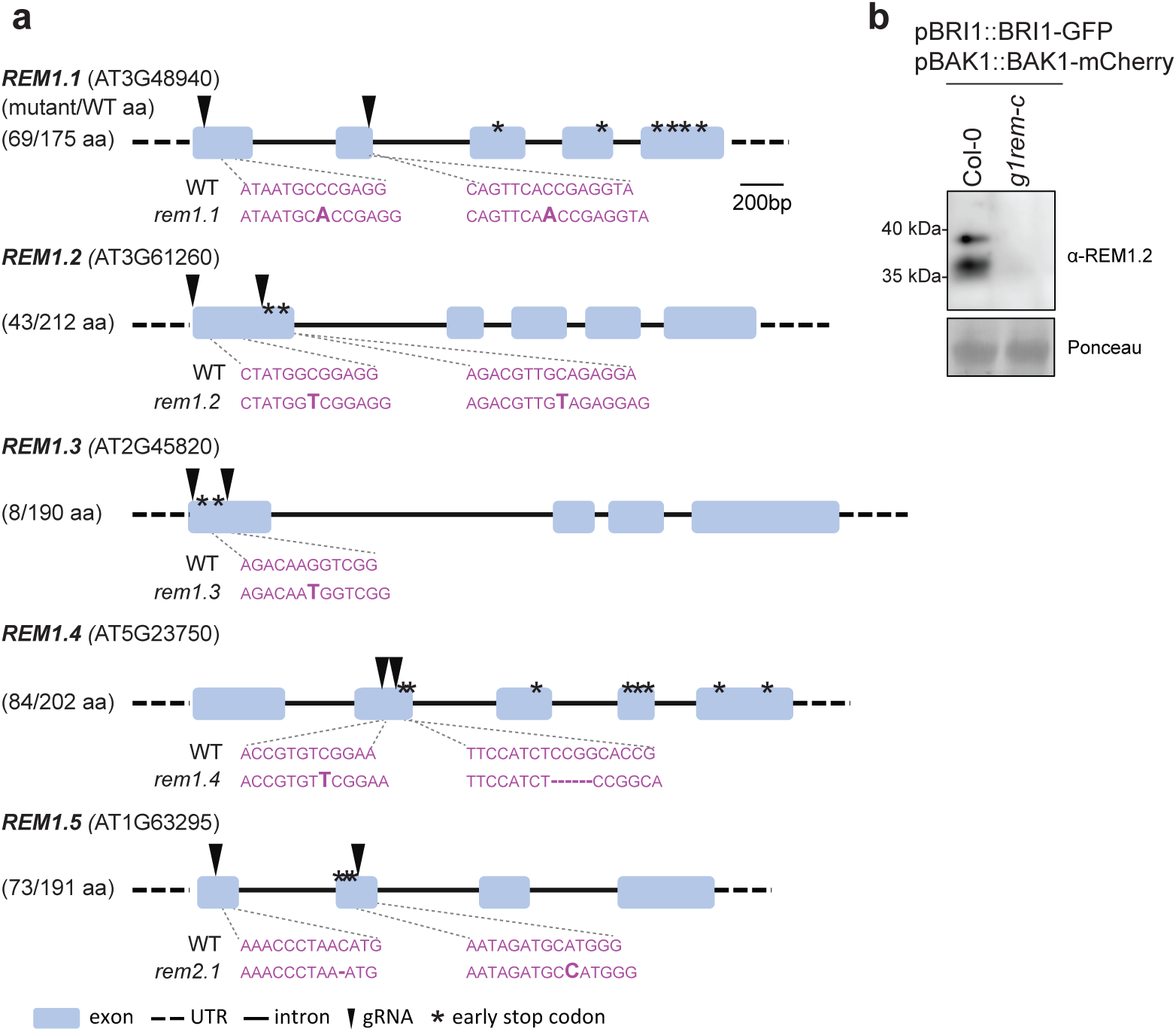
Group 1 *REMs* CRISPR mutant expressing BRI1-GFP and BAK1-mCherry. **a**, Representation of group 1 *REMORIN* genes loci. Black arrows indicate the position targeted by gRNAs. Sequences of *REM* genes in WT and *g1rem-c* are shown in pink, editing events are shown in bold, asterisk indicates early stop codons caused by editing events. Amino acid (aa) numbers indicate the protein and protein fragment expected to be produced WT and *g1rem-c* plants. **b**, Western blot analysis of REM1.2 protein accumulation in Col-0/pBRI1::BRI1-GFP/pBAK1-mCherry and *g1rem-c*/pBRI1::BRI1-GFP/pBAK1-mCherry. The membrane was probed with anti-REM1.2 and stained with Ponceau to assess protein loading.

**Extended Data Figure 11.**
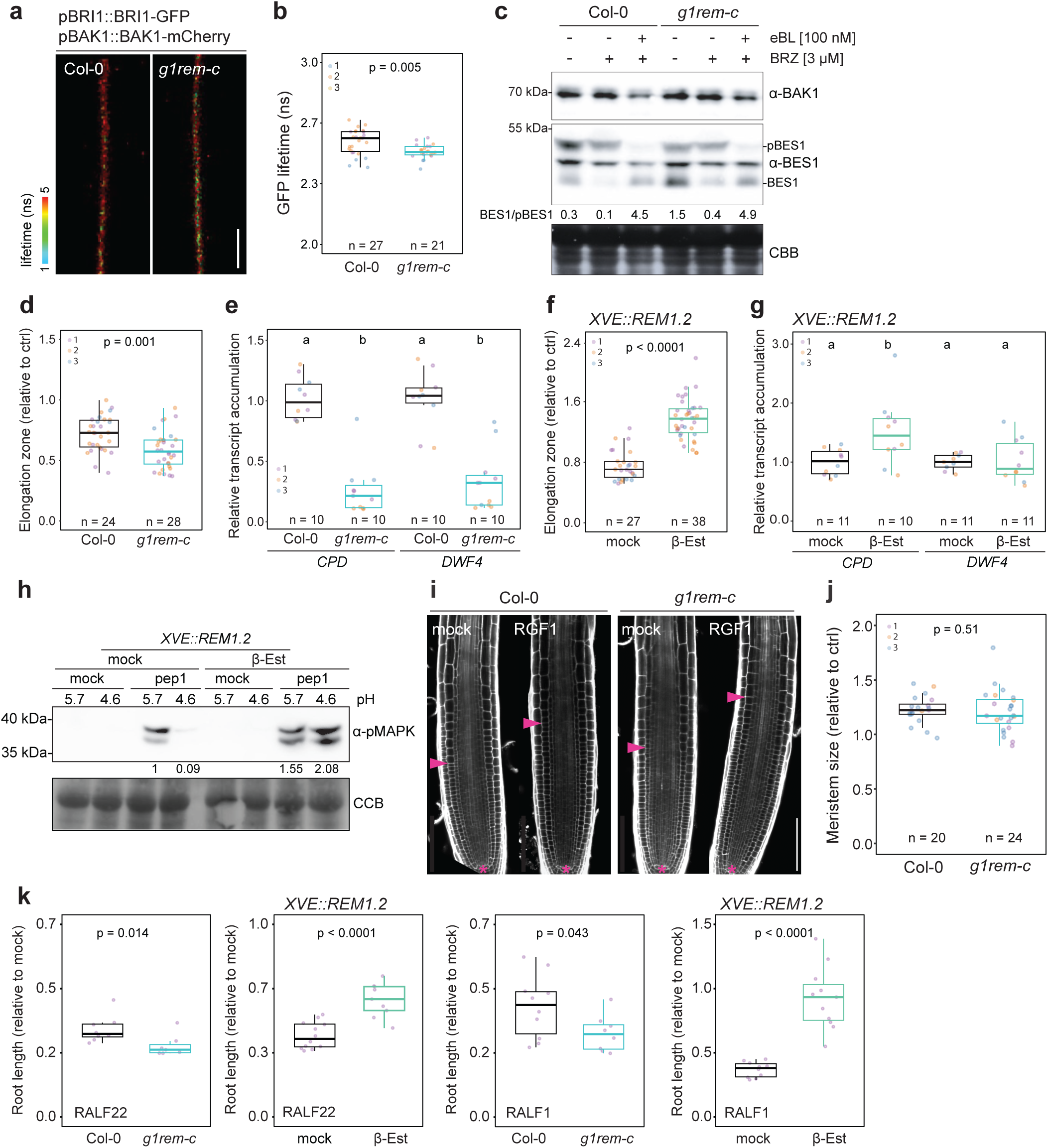
Phenotypic characterization of group 1 *REMs* loss and gain of function mutants. **a**, Fluorescence lifetime imaging microscopy (FLIM) of BRI1-GFP in Col-0/pBRI1::BRI1-GFP/pBAK1-mCherry and *g1rem-c*/pBRI1::BRI1-GFP/pBAK1-mCherry. Scale bar indicates 5 µm. b, Quantification of BRI1-GFP lifetime, data points correspond to individual (n) cells, colors indicate independent experiments. P value reports Mann-Whitney statistical test. **c**, Western blot analysis of BES1 in 12-day-old seedlings in presence and absence of brassinazole (BRZ, an inhibitor of BR synthesis), and eBL. Membranes were probed with anti-BES1, anti-BAK1 antibodies and stained with CBB to assess protein loading. The chemiluminescence intensity ratio of the bands corresponding to BES1 and phosphorylated BES1 (pBES1) are indicated. **d** and **f**, analysis of the effect of exogenous eBL treatment on the length of the elongation zone in 5-day-old seedling roots. Data points correspond to individual (n) roots, colors indicate independent experiments. P values report the results of Mann-Whitney tests. **e** and **g**, Analysis of *CPD* and *DWF4* relative transcript accumulation by RT-qPCR. Data points correspond to independent (n) biological replicates, each corresponding to three 12-day-old seedlings pooled, colors indicate independent experiments. Conditions which do not share a letter are statistically different in Kruskal-Wallis test (p < 0.05). **h**, Western blot analysis of MAPK phosphorylation after pep1 (1 µM) treatment for 5 min in liquid ½ MS media adjusted to pH 5.7 or pH 4.6. Membranes were probed with anti-pERK42/44 (pMAPK) and stained with CBB to assess protein loading. Numbers indicate chemiluminescence intensity ratio of pMAPK normalized to the corresponding CBB intensity and relative to the condition pep1 at pH 5.7. **i**, Representative confocal microscopy images of 5-day-old seedling roots treated with 20 nM RGF1 or corresponding mock control for 24h and stained with propidium iodide. Asterisks indicate quiescent center, arrows mark the end of the meristem, scale bar indicates 100 µm. **j**, Quantification of the effect of RFG1 treatment on the size of the meristem in 5-day-old seedling roots. Data points correspond to individual (n) meristems normalized to mock control conditions, colors indicate independent experiments. P value reports Kruskal–Wallis test. **k**, Analysis of RALF22 and RALF1-induced root growth inhibition. Four-day-old seedlings were treated with 1 µM RALF1 and RAFL22 for 3 days. Data points correspond to individual roots. P values report the results of Mann-Whitney.

**Extended Data Figure 12.**
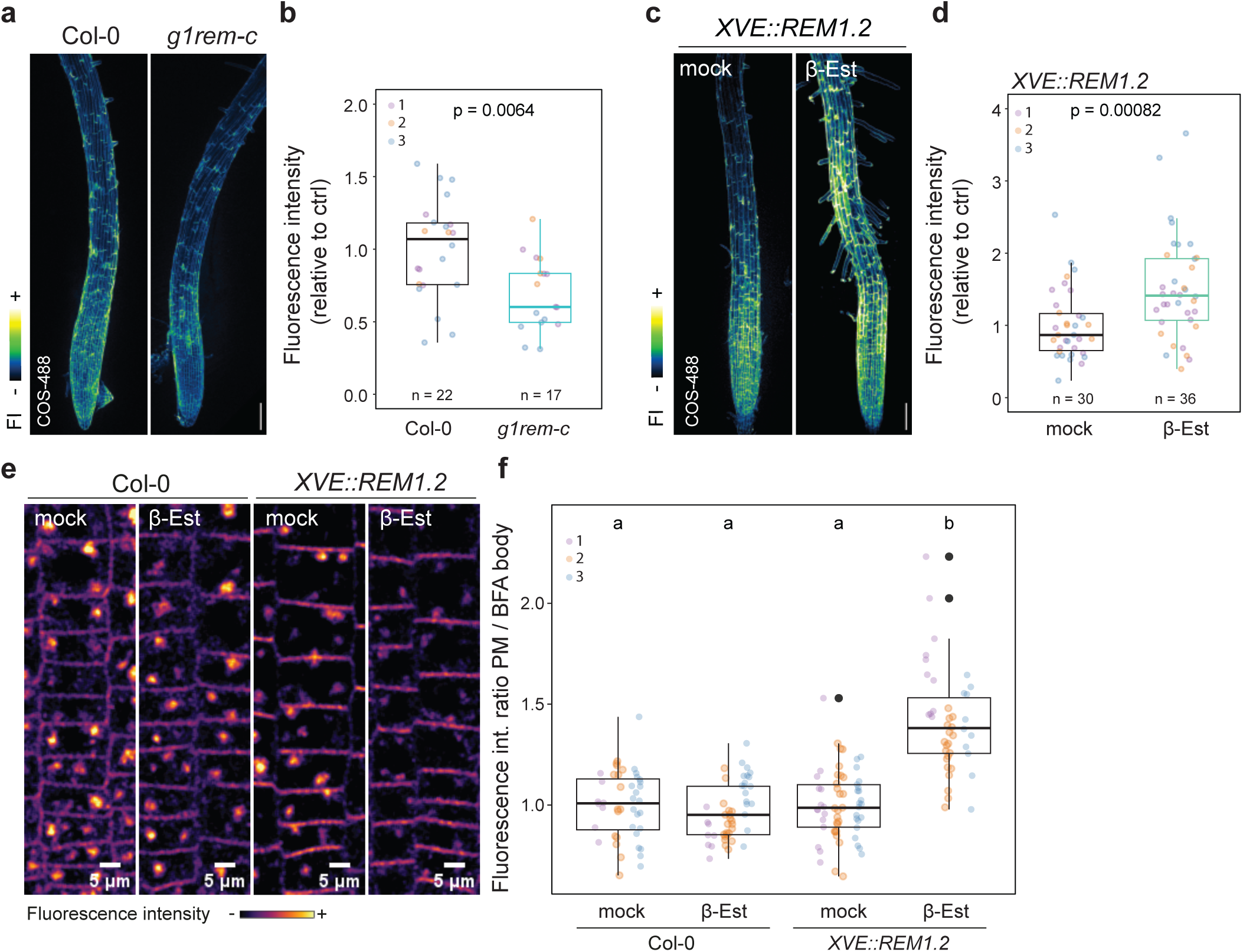
Additional phenotypic characterization of group 1 *REMs* loss and gain of function mutants. **a** and **c**, Confocal microscopy z-projection images of 5-day-old seedling roots stained with chitosan oligosaccharide (COS) coupled with Alexa-488 (COS-488) probing de-esterified pectin oligomers in the cell wall. Scale bar indicates 100 μM. **b** and **d**, Quantification of COS-488 fluorescence intensity. Data points correspond to the average fluorescence intensity in individual (n) root normalized to the control conditions (WT, in **b**) and (mock EtOH treatment in **d**), colors indicate independent experiments. P values report Mann-Whitney tests. **e**, Representative confocal microscopy images of 5-day-old seedling roots treated with Brefeldin A (inhibitor of intracellular vesicular trafficking relying on ARF-GEFs containing Brefeldin A-sensitive SEC7 domains) at 25 µM for 30 min and with the lipophilic fluorescent dye FM4-64 at 1 µM for 5 min. f, analysis of endocytic fluxes based on the ratio of FM4-64 fluorescence at the plasma membrane (PM) and in BFA bodies. Data points correspond to measurements obtained from individual regions of interest (ROI), with 2-4 cells per ROI and 2-3 ROI per root. Data points colors indicate independent experiments. Conditions which do not share a letter are statistically different in pairwise Wilcoxon tests with Bonferroni p-value adjustment (p < 0.001).

**Extended Data Figure 13.**
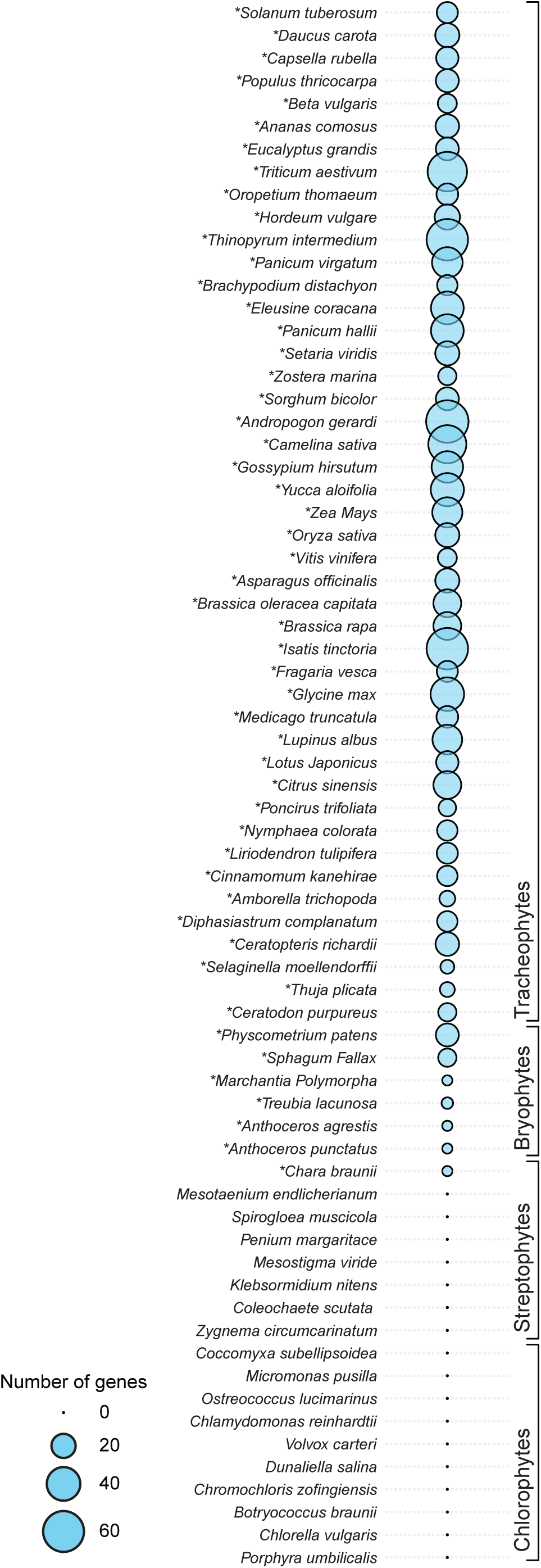
Analysis of the occurrence of REMs in the green lineage. Circles size indicates the number of REM genes in each species. REM genes, transcripts and/or protein occurrence were analyzed by BLAST analyses, PFAMs and INTERPRO domain search. Asterisks indicate species with complex body structure.

**Extended Data Figure 14.**
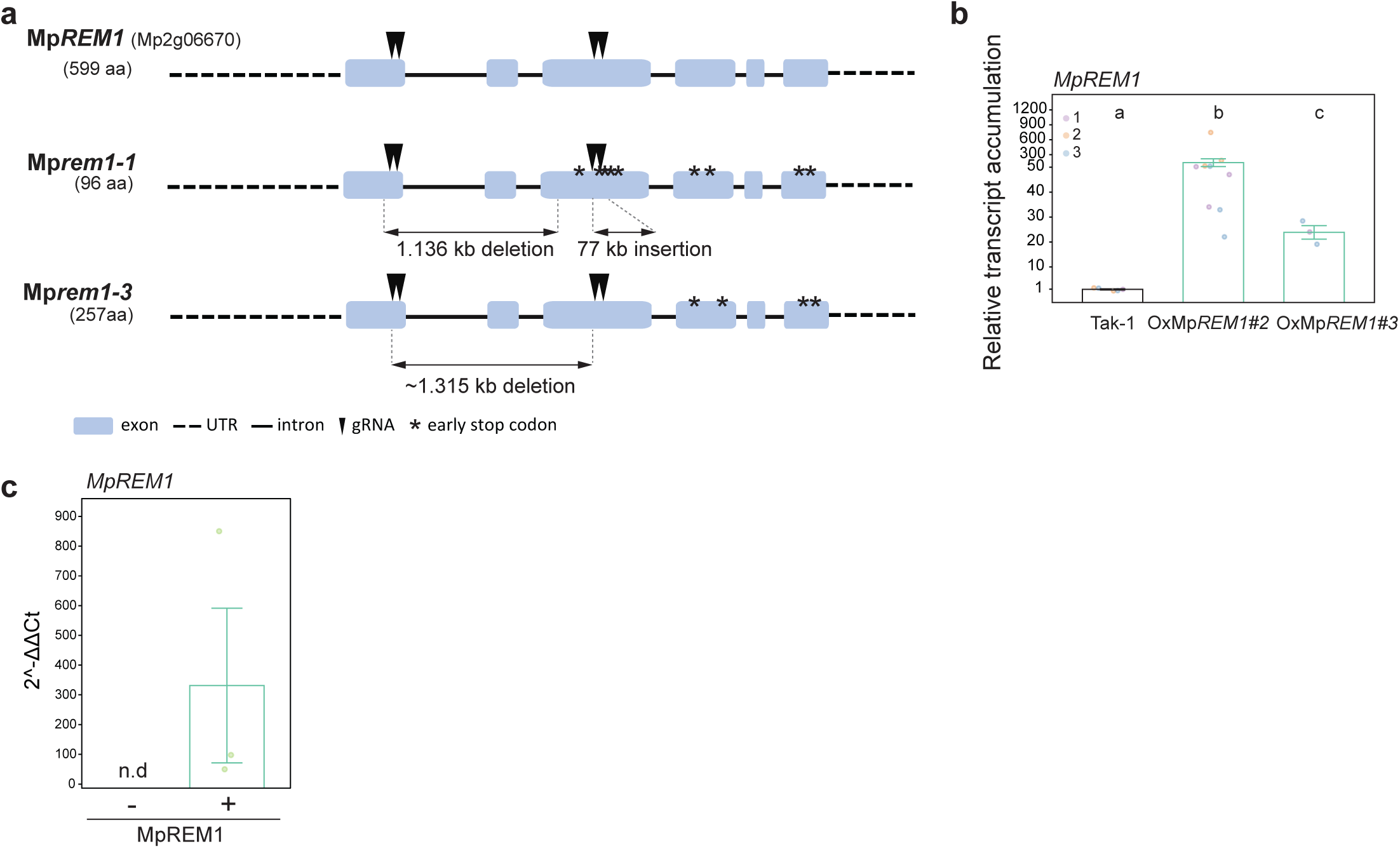
Analysis of *MpREM1* CRISPR knock-out and overexpression lines. **a**, Representation of Mp*REM1* gene locus. Black arrows indicate the position targeted by gRNAs. The nature of the editing events is indicated, asterisks mark the corresponding early stop codons generated. Amino acid (aa) numbers indicate the protein and protein fragments expected to be produced by WT and *MpRem1* plants. **b**, Analysis of Mp*REM1* transcript accumulation in Tak-1, OxMp*REM1-2* and OxMp*REM1-3* transformations. Data points represent relative transcript accumulation in independent biological replicates, colors indicate independent experiments. **c**, Analysis of pUB10::HA-Mp*REM1* transcript accumulation in *Nicotiana benthamiana* leaves. Data points represent independent replicates in infiltrated leaves.

**Extended Data Figure 15.**
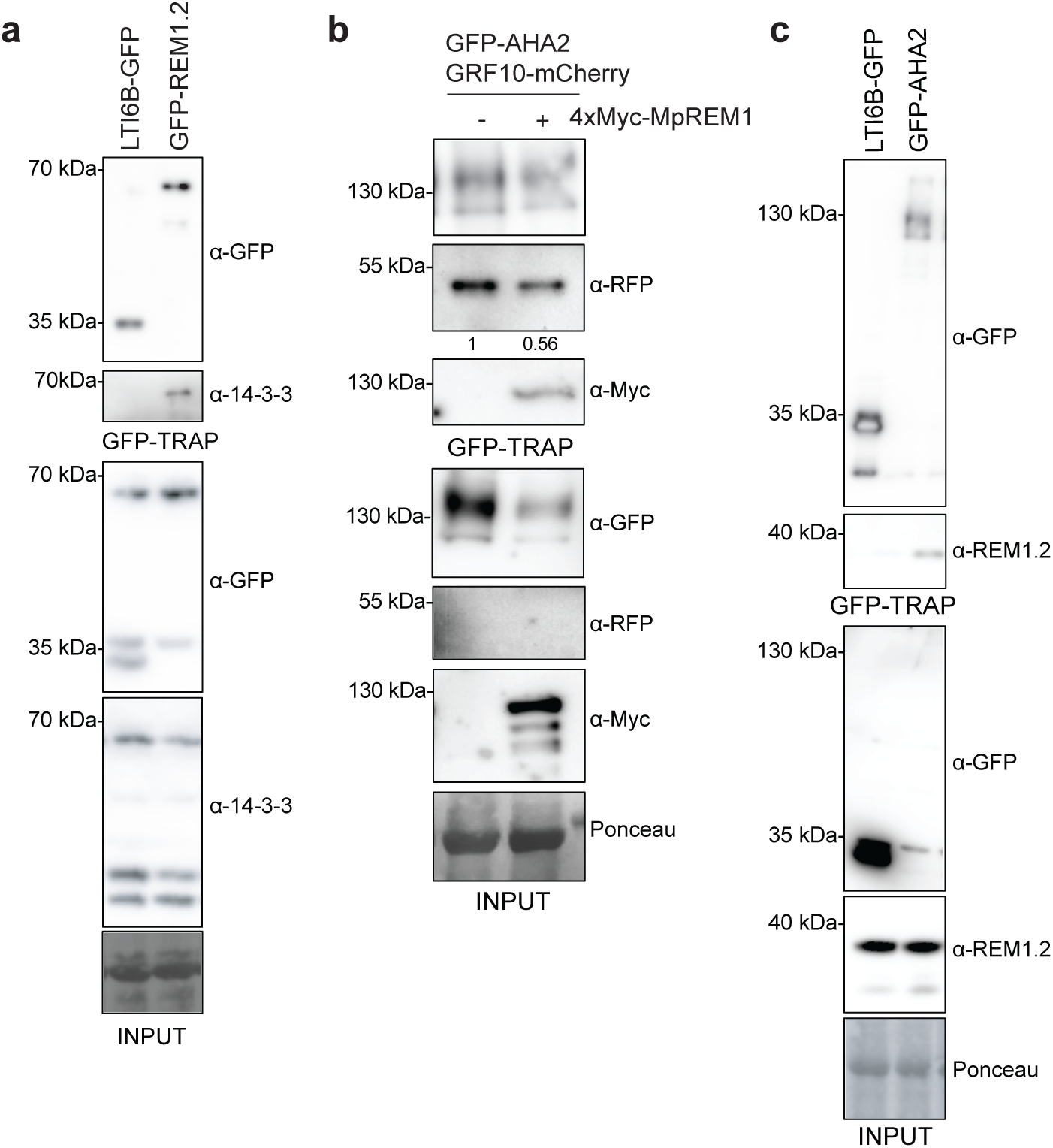
Co-immunoprecipitation experiments. **a**, Immunoprecipitation of p35S::LTI6B-GFP or pUb10::GFP-REM1.2 transiently expressed in *Nicotiana benthamiana leaves.* Membranes were probed with anti-14-3-3 or anti-GFP and stained with Ponceau to assess protein loading. The experiment was repeated thrice with the same results. **b**, Immunoprecipitation experiments of pUb10::GFP-AHA2 transiently co-expressed with pUb10::GRF10-mCherry and p35S::4xMyc-mScarlet-MpREM1 in *Nicotiana benthamiana* leaves treated with 1 μM FC for 1 h. Membranes were probed with anti-RFP (recognizing mCherry), or anti-Myc, or anti-GFP antibodies and stained with Ponceau to assess protein loading. Similar observations were made in at least two independent experiments. **c**, Immunoprecipitation experiments using Col-0/p35S::LTI6B-GFP or *aha2-4*/pAHA2::GFP-AHA2 Arabidopsis seedlings. Membranes were probes with anti-GFP and anti-REM1.2 antibodies and stained with Ponceau to assess protein loading. The experiment was repeated three times with the same results.

**Extended Data Table 1 | List of genes co-expressed with *AHA2*.**

**Extended Data Table 2 | List of primers used in this study.**

**Extended Data Table 3 | List of plasmids used in this study.**

## Data availability

Processed and annotated scRNA-seq data are available at the Gene Expression Omnibus (GSE152766).

## Material and Methods

### Plant materials and growth conditions

*Arabidopsis thaliana* ecotype Col-0 and *Marchantia polymorpha* ecotype Takaragaike-1 (Tak-1, male) were used as wild-type (WT) control and for the generation of stable transgenic lines. The Col-0/p35S::XVE::REM1.2^19^, *rem1.2*/pREM1.2::GFP-REM1.2^19^, Col-0/PM-Apo-Acidin4^11^, Col-0/pBRI1::BRI1-GFP/pBAK1::BAK1-Cherry^53^,*rem1-2*/pREM1.2::mScarlet-AtREM1.2/AtFH6-GFP^16^ and *aha2-4*^54^ mutant lines were previously described. *g1rem-c* was obtained by transforming WT Col-0 with pICSL4723 (*pRPS5a:Cas9i_REM_G1_gRNA1–10_Fast-Red*) (Extended Data Table 3) via floral dipping. Transformants were selected based on seed coat fluorescence, sequenced and T3 Cas9i-negative homozygous *REMs* quintuple mutant was selected for functional analyses. *g1rem-c*/pREM1.2::REM1.2 complementation lines were generated by transforming *g1rem-c* with pREM1.2::REM1.2 (Extended Data Table 3) via floral dipping. The transformants were selected on hygromycin (25 mg/L), and analyzed by western blotting using anti-REM1.2. Col-0/pBRI1::BRI1-GFP/pBAK1::BAK1-Cherry^53^ was transformed with pICSL4723(*pRPS5a:Cas9i_REM_G1_gRNA1–10_Fast-Red*) (Extended Data Table 3) via floral dipping to obtained *g1rem-c*/pBRI1::BRI1-GFP/pBAK1::BAK1-Cherry. Transformants were selected based on seed coat fluorescence, sequenced and T3 Cas9i-negative homozygous *REMs* quintuple mutant was selected for functional analyses. The aha2-4/pAHA2::GFP-AHA2 was generated by transforming *aha2-4*^54^ with pAHA2::GFP-AHA2 (Extended Data Table 3). Mp*rem1* CRISPR mutants were obtained by transforming Marchantia following the previously described cut-thalli transformation method^55^, and transformants were selected on chlorsulfuron selection media, genotyped and sequenced. Mp*rem1-3*/p35S::GFP-MpREM1 was obtained by transforming Mp*rem1-3* with p35S::GFP-MpREM1 (Extended Data Table 3) by cut-thalli transformation method^55^, transformants were selected hygromycin selection media (25 mg/L). Arabidopsis seeds were surface sterilized using 0.1 % Tween 20 in 70 % EtOH for 10 min, following 70 % EtOH for 10 min and 100% EtOH for 1min. Seeds were stratified for 2 days in the dark at 4 °C and grown on half Murashige and Skoog (MS) media supplemented with vitamins, 1 % sucrose and 0.8 % agar at 22 °C and a 16-h light photoperiod. *Nicotiana benthamiana* plants were cultivated under controlled conditions with a 16-hour photoperiod at 25 °C. *Marchantia* plants were grown on 1 % agar half-strength Gamborg B5 (GB5) medium pH 5.5, at 22 °C and a 16-h light photoperiod.

### Molecular cloning, plant transformation and CRISPR-Cas9-mediated mutagenesis

gRNAs for CRISPR Cas-mediated editing were designed using CHOPCHOP (https://chopchop.cbu.uib.no/). Ten gRNAs (Extended Data Table 2) for the multiplexed gene editing of *REMORINs* in *Arabidopsis thaliana* were assembled using Golden Gate cloning^56^ into the BsaI sites of pICH47761, pICH47772, pICH47781, pICH47791, pICH47732, pICH47742 and pICH47751, subsequently assembled into the BpiI sites of pAGM8067 and pAGM8043 which were in turn assembled with the intronic version of Cas9^57^ and the Fast-Red selection marker ^58^ into the BsaI site of pICSL4723 (Extended Data Table 3). For gene editing in *Marchantia polymorpha*, four gRNAs (Extended Data Table 3) were cloned into the BsaI site of pMpGE_En04, pBC-GE12, pBC-GE23 or pBC-GE34. The four gRNA transcriptional units were then transferred into pMpGE018 binary vector carrying the Cas9^D10A^ (nickase)^59,60^ by LR reaction (Invitrogen). AtREM1.2 coding sequence and promoter region were previously described ^61^. The GRF10 and AHA2 coding sequences were obtained as synthetic fragments from TwistBioscience and inserted into the Golden Gate pUC57_BB03 plasmid^62^ using BpiI. Promoter region of *AHA2* was amplified using Arabidopsis Col-0 genomic DNA (gDNA) as template and the primers listed in the Extended Data Table 2 and subsequently cloned into pUC57_BB03 plasmid^62^. Golden gate modules were assembled via BsaI into the Xpre2-S binary BB21_LIIβ R 1-2 or BB29_LIIβ R 5-6 plasmids^62^ to generate pAtREM1.2::AtREM1.2, pUb10::GFP-AtREM1.2, pAHA2::GFP-AHA2, pUb10::GFP-AHA2 and pUb10::GRF10-mCherry. All plasmids were verified by Sanger sequencing. The binary vectors were transferred into Agrobacterium tumefaciens strain GV3101 for flower dip transformation of Arabidopsis or transient transformation of *Nicotiana benthamiana* leaves. Cas9^D10A^-resistant version of MpREM1 coding sequence was obtained as synthetic fragment from TwistBiosience and cloned into the Golden Gate pUC57_BB03 plasmid^62^ and subcloned via BsaI into the Xpre2-S binary LII_ß_F3-4_BB024 plasmid^62^ to obtain p35S::GFP-MpREM1/Hygro, pUb10::MpREM1 and 35S::4xMyc-MpREM1 (Extended Data Table 3).

### Chemicals and peptides

Fusicoccin (FC, Thermofisher), epibrassinolide (eBL, Sigma Cat#E1641) and β-estradiol (β-est, Sigma-Aldrich) were dissolved in ethanol as a stock of 20 mM, 1 mM and 100 mM respectively. RALF1 (ATTKYISYQSLKRNSVPCSRRGASYYNCQNGAQANPYSRGCSKIARCRS), RALF22 (AQKKYISYGAMRRNSVPCSRRGASYYNCQRGAQANPYSRGCSTITRCRR), RGF1 (DY(SO3H)SNPGHHPP(Hydroxy)RHN) and pep1 (ATKVKAKQRGKEKVSSGRPGQHN) synthetic peptides were obtained from SynPeptide CO., LTD at a purity > 95 %, and dissolved in double distilled water to 1 mM. The chemicals were dissolved to final working concentration of 1 μM Fusicoccin, 1 µM RALF1, 1 µM RALF22, 10 nM Pep1, 20 nM RGF1, 1 nM and 5 nM eBL in liquid ½ MS medium pH 5.7.

### Confocal laser scanning microscopy

Confocal microscopy was performed using a Zeiss LSM880 confocal laser scanning microscope, equipped with a Plan-Apochromat 10x/0.45 M27 objective. PI and mCherry were excited with a DPS 561 nm diode laser and fluorescence collected between 570-642 nm. GFP was excited with a 488-nm laser argon laser and fluorescence was collected between 500-550 nm. To obtain quantitative data, experiments were performed using strictly identical acquisition parameters (e.g. laser power, gain, zoom factor, resolution, and emission wavelengths reception), with detector settings optimized for low background and no pixel saturation. Images were processed and analyzed in Fiji^63^.

### Analysis of relative pHe

For 8-hydroxypyrene-1,3,6-trisulfonic acid trisodium salt (HPTS) staining and imaging 5-day-old Arabidopsis seedlings, grown on ½ MS medium with 1 % sucrose, pH 5.7, were mounted on agar supplemented with 1 mM HPTS, as previously described^1^. Confocal images were captured sequentially using a LEICA SP8 confocal microscope. Excitation wavelengths of 458 nm (for deprotonated HPTS) and 405 nm (for protonated HPTS) were used, and emitted light was collected between 485 and 540 nm. The pH sensor PM-Apo-acidin4 was imaged using Zeiss LSM880 confocal laser scanning microscope. Excitation of mRFP was done at 561 nm and the emitted light was recorded between 580 and 650 nm, and SYFP2 was excited at 488 nm and the emitted light was recorded between 510 and 554 nm. The PM-Apo-acidin4 fluorescence ratio was calculated by dividing the SYFP2 signal by the mRFP signal. Ratiometric images were generated as previously described^1^ and analyzed in Fiji^63^.

### COS-488 staining and imaging

Synthesis of COS-488 was carried out as described before^64^. Chitosan oligosaccharides (Carbosynth OC09272) were dissolved in 100 mM sodium acetate buffer pH 4.9 at 1mg/mL. 16 µL of 10 mg/mL AlexaFluor488 hydroxylamine in DMSO were added to 0.5 mL of COS solution and incubated in dark at 37°C for 2 days while shaking. Arabidopsis seedlings were stained and imaged as previously described^65^. Briefly, Arabidopsis seedlings were stained with 1:1000 COS-488 diluted in H_2_O for 10 min and washed twice with H_2_O. Samples were mounted in H_2_O between coverslips and slides imaged using a Zeiss LSM880 confocal laser scanning microscope, equipped with a Plan-Apochromat 10x/0.45 M27 objective. COS-488 was excited with a 488-nm argon laser and fluorescence was collected between 500-550 nm using 600-800 V gain. Z-projections were processed and analyzed in Fiji^63^.

### Fluorescence lifetime imaging microscopy - Förster resonance energy transfer

Fluorescence lifetime imaging microscopy - Förster resonance energy transfer (FLIM-FRET) experiments were performed on Leica SP8 confocal laser scanning microscope (CLSM) (Leica Microsystems GmbH, Wetzlar, Germany) coupled to FastFLIM PicoQuant system consisting of Sepia Multichannel Picosecond Diode Laser, PicoQuant Timeharp 260, TCSPC Module, and Picosecond Event Timer. Imaging was performed using a 63×/1.20 water immersion objective. FLIM measurements were performed with a 440-nm pulsed laser (LDH-P-C-470, Picoquant GmbH, Berlin, Germany) by scanning regions of 256 µm2 with a ~20 µs pixel dwell time and under 40 MHz pulse rate. The maximal count rate was set to ~1500 cps. Fluorescence emission was detected with an HyD4 SMD from 455 nm to 505 nm by time-correlated single-photon counting using a TimeHarp 260 module (PicoQuant GmbH, Berlin, Germany). Calculations of fluorescence lifetime were performed using the PicoQuant SymPhoTime 64 software following instructions for FLIM-FRET calculation for multi-exponential donors. A two-exponential decay fit was used for GFP. The lifetimes were initially estimated by fitting the data using the Monte Carlo method and then by fitting the data using the maximum likelihood estimation.

### Analysis of single-cell RNA sequencing and proteomic data

Processed and annotated data by ^13^ were downloaded from the Gene Expression Omnibus (GSE152766_Root_Atlas_spliced_unspliced_raw_counts.rds.gz). All analyses were performed using log-normalized single-cell RNA-seq data extracted via the R package Seurat (v5.3.0)^66^. Gene expression values for *AHA2* (AT4G30190) *REMs* genes (AT3G48940, AT3G61260, AT2G45820, AT5G23750) together with pseudotime and lineage identifiers were extracted using FetchData. For each lineage, gene expression was smoothed along pseudotime using LOESS regression, and lineage-specific pseudotime expression trajectories were plotted using ggplot2. For the co-expression analyses, cells were selected based on cell-type annotations^13^. For the epidermis data for both atrichoblast and trichoblast were merged before analysis. For each group, pseudotime values were binned into 100 equal bins. The mean expression of every gene was calculated for each bin, genes with missing values across bind were removed. Pseudotime-binned expression profiles were smoothed using natural spline smoothing (ns, df=8). Pearson correlations were computed between the smoothed expression profiles of each gene. To rank transcript abundance, the percentile scores of *AHA1* (AT2G18960), *AHA2* (AT4G30190) *REM1.2* (AT3G61260) and *REM1.3* (AT2G45820) expression were calculated on averaged log-normalized expression values across all cells. Similarly, percentile values, relative to the full Arabidopsis proteome, were calculated based on log10-transformed protein abundance (iBAQ) values from the ATHENA proteomics data set^52^. The following R packages were used for data analyses; dplyr, tidyr, Matrix, ggplot2, ggrepel and future.

### Analysis of Arabidopsis root developmental transitions

To analyze root developmental transition, Arabidopsis seedlings were stained with 10 μg/mL propidium iodide (PI) dissolved in water for 10 min, then washed twice in water. Samples were mounted in H2O on coverslips and analyzed at 600-640 nm for propidium iodide staining using Zeiss LSM880 confocal laser scanning microscope. The length of root meristems was obtained by measuring length of the cortex cells from the cell proximal to the quiescent center to thirst rapid elongated cell. The initiation of differentiation was obtained by measuring the length from cell proximal to the QC to the first root hair bulging. Measurements were done using Fiji^63^.

### Analysis of root hair length

Five-day-old Arabidopsis seedlings, grown on ½ MS medium with 1 % sucrose, pH 5.7, were imaged using stereo-microscope (ZEISS Axio Zoom.V16). Root hair length was measured using Fiji^63^.

### Transient transformation of *Nicotiana benthamiana* leaves

For transient protein expression in *Nicotiana benthamiana*, leaves of 4-week-old plants were infiltrated with Agrobacterium tumefaciens strain GV3101 carrying plasmids the corresponding constructs as indicated in figure legends. *N. benthamiana* leaves were collected two days post-infiltration.

### Microsomes preparations

Seedlings were grown on ½ MS medium for 4 days and transferred to liquid ½ MS medium for two weeks. Marchantia plants were grown in ½ GB5 media for 20 days. Samples were isolated and frozen in liquid nitrogen. Proteins were isolated in homogenization buffer (50 mM Tris-HCl, pH 7.5, 150 mM NaCl; 10% glycerol, 2 mM EDTA, 5 mM DTT, 1 % protease inhibitor cocktail, 1 % IGEPAL, 1 mM NaF, 1 mM PMSF). Protein samples were clarified by centrifugation, 13 000 x g for 20 min at 4 °C. Microsomal fractions were collected from supernatant by ultracentrifugation at 100 000 x g for 1 hour at 4 °C.

### Measurement of vanadate-sensitive ATPase activity

Vanadate-sensitive ATP hydrolysis was measured in the microsomal fractions following the methods described in ^67,68^. The microsomal fraction (20 µg protein; 0.45 µg/µL) was mixed with an equal volume of ATPase reaction buffer (60 mM MES–Tris, pH 6.5; 6 mM MgSO₄; 200 mM KNO₃; 100 mM KCl; 1 mM ammonium molybdate; 10 µg/mL oligomycin; 0.1 % Triton X-100; 1 mM PMSF; and 1 % protease inhibitor cocktail (Sigma-Aldrich), with or without 2 µL of 10 mM sodium orthovanadate. The reaction was initiated by adding 10 µL of 20 mM ATP (Roth, Art. No. HN35.1) and incubated at 24 °C for 30 min. The reaction was terminated by adding 1 mL of stop solution (1.3 % [w/v] SDS, 0.25 % [w/v] sodium molybdate, and 0.3 N H₂SO₄) and 50 µl ANSA solution (0.125 % (w/v) 1-amino-2-naphthol-4-sulfonic acid (ANSA), 15 % (w/v) NaHSO3, 1 % (w/v) Na2SO4). Absorbance was measured at 750 nm using a spectrophotometer. Inorganic phosphate (Pi) content (nmol) was determined from a standard curve generated with known Pi concentrations using the equation:

Absorbance₇₅₀ = (slope × Pi content [nmol]) + y-intercept.

Vanadate-sensitive ATP hydrolytic activity (nmol Pi h⁻¹ mg⁻¹ protein) was calculated as: ([Pi]₋vanadate − [Pi]₊vanadate) / (incubation time) / (protein amount).

### RNA isolation and quantitative RT-PCR

RNA isolation, cDNA synthesis and quantitative real-time PCR (qPCR) was performed similarly as previously described in^35^. Samples were collected from 12-day-old Arabidopsis seedlings grown 4 days in solid ½ MS 1% sucrose pH 5.7 and 8 days in liquid ½ MS 1 % sucrose pH 5.7 or from 3-week-old Marchantia thallus grown on ½ GB5, pH 5.5 medium. Each sample was transferred to 2 mL tubes containing 2 mm glass beads and flash-frozen in liquid nitrogen. Samples were grinded while frozen for 1.5 minutes at 1.500 rpm using BioRad TissueLyser. RNA was extracted using TRIzol™ Reagent (ThermoFischer scientific). 200 ng of RNA were used for first-strand cDNA synthesis which was performed using RevertAid first strand cDNA synthesis kit (ThermoFischer scientific) and oligo(dT)18 according to the manufacturer’s instructions. cDNA was amplified in triplicate by quantitative PCR by using PowerUp™ SYBR™ Green Master Mix (ThermoFischer scientific) and a 7500 Fast real-time PCR detection system. Primers used to assess transcript accumulation are listed in (Extended Data Table 2) Expression levels were normalized to those for *UBOX*, using the comparative Ct method (2-ΔΔCt).

### Co-immunoprecipitation

Immunoprecipitations were performed as previously described^69^. For experiments in Arabidopsis, 20–30 seedlings per plate were grown in wells of a 6-well plate for 2 weeks, transferred to 2 mM MES-KOH, pH 5.7, and incubated overnight. Seedlings were then frozen in liquid nitrogen and subjected to protein extraction. For transient expression in *N. benthamiana*, leaves of four-week-old plants were infiltrated with *Agrobacterium tumefaciens* strain GV3101 carrying the constructs described in the figures. GV3101 harboring the p19 RNA-silencing suppressor was co-infiltrated in all cases. Leaf tissues were collected 2 days post-infiltration, frozen in liquid nitrogen and subjected to protein extraction. Proteins were isolated in 50 mM Tris-HCl pH 7.5, 150 mM NaCl, 10 % glycerol, 5 mM dithiothreitol, 1 % protease inhibitor cocktail (Sigma-Aldrich), 2 mM Na2MoO4, 2.5 mM NaF, 1.5 mM activated Na3VO4, 1 mM phenylmethanesulfonyl fluoride, and 0.5% IGEPAL. For immunoprecipitations, GFP-Trap agarose beads (ChromoTek) were used and incubated with the crude extract for 3 to 4 h at 4 °C. Subsequently, beads were washed three times with wash buffer (50 mM Tris-HCl pH 7.5, 150 mM NaCl, 1 mM phenylmethanesulfonyl fluoride, 0,1 % IGEPAL) before adding Laemmli sample buffer and incubating for 10 min at 95°C. Analysis was carried out by SDS-PAGE and immunoblotting.

### Mobility shift-based estimation of BES1 phosphorylation status

Epibrassinolide (eBL)-induced BES1 dephosphorylation assays were performed as previously described^70^ seedlings were germinated on half MS-agar supplemented with 1 % sucrose for five days before transplanting to 24-well plates (two seedlings per well) containing liquid half MS supplemented with 1 % sucrose. One day before the assays, the medium was replaced with fresh medium. Twelve-day-old seedlings were treated with eBL or ethanol (mock) at the indicated concentrations for 10 min. Samples were blot-dried, transferred to 2 mL tubes containing 2 mm glass beads and flash-frozen in liquid nitrogen and store at −80 ◦C before protein extraction and immunoblotting.

### Immunoblotting

Protein samples were separated in 8-10% bis-acrylamide gels at 150 V for approximately 2 hours and transferred into activated PVDF membranes at 100 V for 90 min. Immunoblotting was performed with antibodies diluted in blocking solution (5 % milk in TBS with 1 % [v/v] Tween-20). Antibodies used in this study were α-REM1.2 (1:3000)(von Arx et al., 2025), α-GFP (1:5000, sc-9996, Santa Cruz), α-BES1 (1:2000)^70^, α-BAK1 (1:5000)^71^, α-pErk44/42 (1:5000, 137F5 #4695, Cell Signaling Technology), α-14-3-3 (1:2000, Agrisera AS122119). Blots were developed with Pierce ECL/ECL Femto Western Blotting Substrate (Thermo Scientific). The following secondary antibody was used: anti-rabbit IgG (whole molecule)–HRP 1:10000 (A0545, Sigma).

### Statistical analysis

The number of independent experiments, the number of individual cells or roots analyzed per condition and collected across these experiments are indicated in each figure legend. The statistical tests used are reported in the figure legends and have been performed using R.

## References

1. Barbez, E., Dünser, K., Gaidora, A., Lendl, T. & Busch, W. Auxin steers root cell expansion via apoplastic pH regulation in Arabidopsis thaliana. Proc Natl Acad Sci U S A 114, E4884–E4893 (2017).

2. Martinière, A. et al. Uncovering pH at both sides of the root plasma membrane interface using noninvasive imaging. Proc Natl Acad Sci U S A 115, 6488–6493 (2018).

3. Oginuma, M. et al. Intracellular pH controls WNT downstream of glycolysis in amniote embryos. Nature 584, 98–101 (2020).

4. Liu, Y. et al. Intracellular pH dynamics regulates intestinal stem cell lineage specification. Nat Commun 14, (2023).

5. Peters, W. S. & Felle, H. H. The Correlation of Profiles of Surface pH and Elongation Growth in Maize Roots. Plant Physiol 121, 905–912 (1999).

6. Xu, F. & Yu, F. Sensing and regulation of plant extracellular pH. Trends Plant Sci 28, 1422–1437 (2023).

7. Casey, J. R., Grinstein, S. & Orlowski, J. Sensors and regulators of intracellular pH. Nat Rev Mol Cell Biol 11, 50–61 (2010).

8. Tsai, H. H. & Schmidt, W. The enigma of environmental pH sensing in plants. Nat Plants 7, 106–115 (2021).

9. Xhelilaj, K., Fuglsang, A. T., Gronnier, J. & Palmgren, M. Molecular tipping points in plant cell surface H+ homeostasis and signalling. Quantitative Plant Biology 6, e28 (2025).

10. Miao, R., Russinova, E. & Rodriguez, P. L. Tripartite hormonal regulation of plasma membrane H+-ATPase activity. Trends Plant Sci 27, 588–600 (2022).

11. Moreau, H., Gaillard, I. & Paris, N. Genetically encoded fluorescent sensors adapted to acidic pH highlight subdomains within the plant cell apoplast. J Exp Bot 73, 6744–6757 (2022).

12. Palmgren, M. G. PLANT PLASMA MEMBRANE H+-ATPases: Powerhouses for Nutrient Uptake. Annu Rev Plant Physiol Plant Mol Biol 52, 817–845 (2001).

13. Shahan, R. et al. A single-cell Arabidopsis root atlas reveals developmental trajectories in wild-type and cell identity mutants. Dev Cell 57, 543–560.e9 (2022).

14. Raffaele, S., Mongrand, S., Gamas, P., Niebel, A. & Ott, T. Genome-wide annotation of remorins, a plant-specific protein family: Evolutionary and functional perspectives. Plant Physiol 145, 593–600 (2007).

15. Gronnier, J. et al. Structural basis for plant plasma membrane protein dynamics and organization into functional nanodomains. Elife 6, (2017).

16. Ma, Z. et al. Membrane nanodomains modulate formin condensation for actin remodeling in Arabidopsis innate immune responses. Plant Cell 34, 374–394 (2022).

17. Su, C. et al. Stabilization of membrane topologies by proteinaceous remorin scaffolds. Nature Communications 2023 14:1 14, 323- (2023).

18. Jolivet, M.-D. et al. Interdependence of plasma membrane nanoscale dynamics of a kinase and its cognate substrate underlies Arabidopsis response to viral infection. Elife 12, (2025).

19. Huang, D. et al. Salicylic acid-mediated plasmodesmal closure via Remorin-dependent lipid organization. Proc Natl Acad Sci U S A 116, 21274–21284 (2019).

20. Cantley, L. C., Resh, M. D. & Guidotti, G. Vanadate inhibits the red cell (Na+, K+) ATPase from the cytoplasmic side. Nature 1978 272:5653 272, 552–554 (1978).

21. Palmgren, M. G. & Christensen, G. Functional comparisons between plant plasma membrane H(+)-ATPase isoforms expressed in yeast. Journal of Biological Chemistry 269, 3027–3033 (1994).

22. Pacifici, E., Di Mambro, R., Dello Ioio, R., Costantino, P. & Sabatini, S. Acidic cell elongation drives cell differentiation in the Arabidopsis root. EMBO J 37, EMBJ201899134- (2018).

23. Jahn, T. et al. The 14-3-3 protein interacts directly with the C-terminal region of the plant plasma membrane H(+)-ATPase. Plant Cell 9, 1805–1814 (1997).

24. Fuglsang, A. T. & Palmgren, M. Proton and calcium pumping P-type ATPases and their regulation of plant responses to the environment. Plant Physiol 187, 1856–1875 (2021).

25. Hoffmann, R. D. et al. Roles of plasma membrane proton ATPases AHA2 and AHA7 in normal growth of roots and root hairs in Arabidopsis thaliana. Physiol Plant 166, 848–861 (2019).

26. Lin, C. Y. et al. Pathways involved in vanadate-induced root hair formation in Arabidopsis. Physiol Plant 153, 137–148 (2015).

27. Stéger, A. & Palmgren, M. Root Hair Growth from the PH Point of View. (2022) doi:10.3389/fpls.2022.949672.

28. Zhai, K. et al. A Phosphorelay Circuit Drives Extracellular Alkalinization in Plant Receptor Kinase Signaling. bioRxiv 2025.08.16.670655 (2025) doi:10.1101/2025.08.16.670655.

29. Liu, L. et al. Extracellular pH sensing by plant cell-surface peptide-receptor complexes. Cell 185, 3341–3355.e13 (2022).

30. Diaz-Ardila, H. N., Gujas, B., Wang, Q., Moret, B. & Hardtke, C. S. pH-dependent CLE peptide perception permits phloem differentiation in Arabidopsis roots. Current Biology 33, 597–605.e3 (2023).

31. Sun, Y. et al. Structure reveals that BAK1 as a co-receptor recognizes the BRI1-bound brassinolide. Cell Research 2013 23:*11* 23, 1326–1329 (2013).

32. González-García, M. P. et al. Brassinosteroids control meristem size by promoting cell cycle progression in Arabidopsis roots. Development 138, 849–859 (2011).

33. Hacham, Y. et al. Brassinosteroid perception in the epidermis controls root meristem size. Development 138, 839–848 (2011).

34. Yin, Y. et al. BES1 accumulates in the nucleus in response to brassinosteroids to regulate gene expression and promote stem elongation. Cell 109, 181–191 (2002).

35. Albrecht, C. et al. Brassinosteroids inhibit pathogen-associated molecular pattern-triggered immune signaling independent of the receptor kinase BAK1. Proc Natl Acad Sci U S A 109, 303–308 (2012).

36. Tang, J. et al. Structural basis for recognition of an endogenous peptide by the plant receptor kinase PEPR1. Cell Research 2015 25:1 25, 110–120 (2014).

37. Jing, Y. et al. Danger-Associated Peptides Interact with PIN-Dependent Local Auxin Distribution to Inhibit Root Growth in Arabidopsis. Plant Cell 31, 1767–1787 (2019).

38. Matsuzaki, Y., Ogawa-Ohnishi, M., Mori, A. & Matsubayashi, Y. Secreted peptide signals required for maintenance of root stem cell niche in Arabidopsis. Science 329, 1065–1067 (2010).

39. Haruta, M., Sabat, G., Stecker, K., Minkoff, B. B. & Sussman, M. R. A peptide hormone and its receptor protein kinase regulate plant cell expansion. Science 343, 408–411 (2014).

40. Schoenaers, S. et al. Rapid alkalinization factor 22 has a structural and signalling role in root hair cell wall assembly. Nat Plants 10, 494–511 (2024).

41. Moussu, S. et al. Plant cell wall patterning and expansion mediated by protein-peptide-polysaccharide interaction. Science (1979) 382, 719–725 (2023).

42. Sénéchal, F. et al. Structural and dynamical characterization of the pH-dependence of the pectin methylesterase–pectin methylesterase inhibitor complex. Journal of Biological Chemistry 292, 21538–21547 (2017).

43. Sakai, H. et al. Increases in intracellular pH facilitate endocytosis and decrease availability of voltage-gated proton channels in osteoclasts and microglia. Journal of Physiology 591, 5851–5866 (2013).

44. Ben-Dov, N. & Korenstein, R. Proton-induced endocytosis is dependent on cell membrane fluidity, lipid-phase order and the membrane resting potential. Biochimica et Biophysica Acta (BBA) - Biomembranes 1828, 2672–2681 (2013).

45. Nishiyama, T. et al. The Chara Genome: Secondary Complexity and Implications for Plant Terrestrialization. Cell 174, 448–464.e24 (2018).

46. de Vries, J. & Archibald, J. M. Plant evolution: landmarks on the path to terrestrial life. New Phytologist 217, 1428–1434 (2018).

47. Harris, B. J. et al. Divergent evolutionary trajectories of bryophytes and tracheophytes from a complex common ancestor of land plants. Nature Ecology & Evolution 2022 6:11 6, 1634–1643 (2022).

48. Bowman, J. L. et al. Insights into Land Plant Evolution Garnered from the Marchantia polymorpha Genome. Cell 171, 287–304.e15 (2017).

49. Fuglsang, A. T. et al. Binding of 14-3-3 protein to the plasma membrane H+-ATPase AHA2 involves the three C-terminal residues Tyr946-Thr-Val and requires phosphorylation of Thr947. Journal of Biological Chemistry 274, 36774–36780 (1999).

50. Li, L., Gallei, M. & Friml, J. Bending to auxin: fast acid growth for tropisms. Trends Plant Sci 27, 440–449 (2022).

51. Lin, W. et al. TMK-based cell-surface auxin signalling activates cell-wall acidification. Nature 2021 599:7884 599, 278–282 (2021).

52. Mergner, J. et al. Mass-spectrometry-based draft of the Arabidopsis proteome. Nature 2020 579:7799 579, 409–414 (2020).

53. Hutten, S. J. et al. Visualization of BRI1 and SERK3/BAK1 Nanoclusters in Arabidopsis Roots. PLoS One 12, e0169905 (2017).

54. Li, L. et al. RALF1 peptide triggers biphasic root growth inhibition upstream of auxin biosynthesis. Proc Natl Acad Sci U S A 119, e2121058119 (2022).

55. Kubota, A., Ishizaki, K., Hosaka, M. & Kohchi, T. Efficient Agrobacterium-mediated transformation of the liverwort Marchantia polymorpha using regenerating thalli. Biosci Biotechnol Biochem 77, 167–172 (2013).

56. Engler, C. et al. A Golden Gate Modular Cloning Toolbox for Plants. ACS Synth Biol 3, 839–843 (2014).

57. Grützner, R. et al. High-efficiency genome editing in plants mediated by a Cas9 gene containing multiple introns. Plant Commun 2, 100135 (2021).

58. Shimada, T. L., Shimada, T. & Hara-Nishimura, I. A rapid and non-destructive screenable marker, FAST, for identifying transformed seeds of Arabidopsis thaliana. Plant J 61, 519–528 (2010).

59. Ran, F. A. et al. Double Nicking by RNA-Guided CRISPR Cas9 for Enhanced Genome Editing Specificity. Cell 154, 1380–1389 (2013).

60. Shen, B. et al. Efficient genome modification by CRISPR-Cas9 nickase with minimal off-target effects. Nat Methods 11, 399–402 (2014).

61. Arx, M. von Xhelilaj, K., Schulz, P., Oven-Krockhaus, S. zur & Gronnier, J. Photochromic reversion enables long-term tracking of single molecules in living plants. bioRxiv 2024.04.10.585335 (2024) doi:10.1101/2024.04.10.585335.

62. Binder, A. et al. A Modular Plasmid Assembly Kit for Multigene Expression, Gene Silencing and Silencing Rescue in Plants. PLoS One 9, e88218 (2014).

63. Schindelin, J. et al. Fiji: an open-source platform for biological-image analysis. Nature Methods 2012 9:*7* 9, 676–682 (2012).

64. Mravec, J. et al. An oligogalacturonide-derived molecular probe demonstrates the dynamics of calcium-mediated pectin complexation in cell walls of tip-growing structures. Plant J 91, 534–546 (2017).

65. Biermann, D. et al. A RALF-brassinosteroid signaling circuit regulates Arabidopsis hypocotyl cell shape. Current Biology 35, 5002–5017.e5 (2025).

66. Hao, Y. et al. Integrated analysis of multimodal single-cell data. Cell 184, 3573–3587.e29 (2021).

67. Oecking, C., Piotrowski, M., Hagemeier, J. & Hagemann, K. Topology and target interaction of the fusicoccin-binding 14-3-3 homologs of Commelina communis. The Plant Journal 12, 441–453 (1997).

68. Okumura, M. & Kinoshita, T. Measurement of ATP Hydrolytic Activity of Plasma Membrane H+-ATPase from <em>Arabidopsis thaliana</em> Leaves. Bio Protoc 6, (2016).

69. Kadota, Y. et al. Quantitative phosphoproteomic analysis reveals common regulatory mechanisms between effector- and PAMP-triggered immunity in plants. New Phytologist 221, 2160–2175 (2019).

70. Perraki, A. et al. Phosphocode-dependent functional dichotomy of a common co-receptor in plant signaling. Nature 561, 248 (2018).

71. Chinchilla, D. et al. A flagellin-induced complex of the receptor FLS2 and BAK1 initiates plant defence. Nature 448, 497–500 (2007).

